# Multi-environment QTL analysis delineates a major locus associated with homoeologous exchanges for water-use efficiency and seed yield in allopolyploid *Brassica napus*

**DOI:** 10.1101/2021.07.08.451711

**Authors:** Harsh Raman, Rosy Raman, Ramethaa Pirathiban, Brett McVittie, Niharika Sharma, Shengyi Liu, Yu Qiu, Anyu Zhu, Andrzej Killian, Brian Cullis, Graham D. Farquhar, Hilary S. Williams, Rosemary White, David Tabah, Andrew Easton, Yuanyuan Zhang

**Affiliations:** NSW Department of Primary Industries, Wagga Wagga Agricultural Institute, Wagga Wagga, NSW 2650, Australia; Centre for Biometrics and Data Science for Sustainable Primary Industries, National Institute for Applied Statistics Research Australia, University of Wollongong NSW 2522, Australia; NSW Department of Primary Industries, Orange Agricultural Institute, ORANGE, NSW 2800, Australia; Oil Crops Research Institute, Chinese Academy of Agricultural Sciences, Wuhan, Hubei, 430062, China; Diversity Arrays Technology P/L, University of Canberra, ACT 2601, Australia; Research School of Biology, Australian National University, Canberra ACT 2601, Australia; CSIRO, Canberra, ACT 2601, Australia; Advanta Seeds Pty Ltd, 268 Anzac Avenue, Toowoomba, QLD 4350, Australia

**Keywords:** water use efficiency, carbon isotope discrimination, drought avoidance, genetic analysis, physiology, yield

## Abstract

- Canola varieties exhibit discernible variation in drought avoidance and drought escape traits, suggesting its adaptation to water-deficit environments. However, the underlying mechanisms are poorly understood.
- A doubled haploid (DH) population was analysed to identify QTL associated with water use efficiency (WUE) related traits. Based on the resequenced parental genome data, we developed sequence-capture based markers for fine mapping. mRNA-Seq was performed to determine the expression of candidate genes underlying QTL for carbon isotope discrimination (Δ^13^C).
- QTL contributing to main and QTL × Environment interaction effects for Δ^13^C and for agronomic WUE were identified. One multi-trait QTL for Δ^13^C, days to flower, plant height and seed yield was identified on chromosome A09, in the vicinity of *ERECTA*. Interestingly, this QTL region was overlapped with a homoeologous exchange event (HE), suggesting its association with the major QTL. Transcriptome analysis revealed several differentially expressed genes between parental lines, including in HE regions.
- This study provides insights into the complexity of WUE related genes in the context of canola adaptation to water-deficit conditions. Our results suggest that alleles for high Δ^13^C contribute positively to canola yield. Genetic and genomic resources developed herein could be utilised to make genetic gains for improving canola WUE.

## Introduction

Drought is the major abiotic stress that reduces the yield potential of various crops, especially in arid and semi-arid regions, of which 89% of regions are prevalent in Oceania (Koohafkan & Stewart, 2008). No doubt the impact of drought stress on crop productivity can be alleviated through irrigation at the ‘critical’ stages of plant development. However, in recent years fresh water, suitable for irrigation, is becoming scarce for crop production, required to meet the demand of a burgeoning human population (Gleick, 2000). Predicted climatic patterns such as debilitating drought and heat-wave episodes and their possible increased frequency further pose a significant threat to crop production (Smith & De Smet, 2012; Mills et al., 2018). The proportion of arable land per capita is also decreasing at a significant rate due to population growth and land degradation (http://www.fao.org/sustainability/). Therefore, improving crop varieties that have high yield potential and utilise water more effectively or require less water could provide a part of the solution to reduce the negative impacts of drought stress and increase productivity and food security (Passioura, 1977; Kijne et al., 2003; Blum, 2009; Bertolino et al., 2019; Leakey et al., 2019).

In nature, to cope with water-deficit conditions, plants have evolved different strategies such as drought escape, drought avoidance and drought tolerance (Levitt, 1980; Ludlow, 1989; Zhu et al., 2016; Rodrigues et al., 2019). Through tiny microscopic pores in the surface of leaves called stomata, plants assimilate CO_2_ for photosynthesis by trading-off water, required for transpiration and other biological processes. This close intimacy between productivity and water use contributes to the adaptation of plants to their growing environments. Therefore, genetic variation in WUE and transpiration efficiency (TE, biomass production/transpirational water loss) that occurs as a result of intentional (via breeding/selection) and unintentional selection in nature provides an opportunity to identify and assemble useful alleles for improving the productivity of various crops.

WUE can be measured at the single leaf level as intrinsic WUE (*i*WUE), defined as the ratio of the photosynthetic CO_2_ assimilation rate (*A*) over transpirational water loss (stomatal conductance, *g*_*sw*_) or as whole-plant vegetative WUE, as the ratio of total dry matter production to total water transpired or as an integrated whole-plant WUE, as the ratio of biomass or seed yield to evapotranspiration (Farquhar & Richards, 1984; Zhengbin et al., 2011; Leakey et al., 2019; Raman et al., 2019). *i*WUE assessments using the gas-exchange method are very challenging to be accurately performed, particularly in the large breeding populations, as WUE is regulated by a myriad of plant development, physiological, biochemical and molecular networks (Moore et al., 2009; Takahashi et al., 2018). Farquhar and Richards (1984) proposed Δ^13^C as a time-integrated surrogate trait for measuring TE both at the single leaf level and at the whole plant level, as C_3_ plants discriminate less against ^13^C during photosynthesis with increased water deficit stress. The negative relationship between WUE and Δ^13^C has been verified in *A. thaliana* (Masle et al., 2005) and in some agricultural crop plants (Farquhar et al., 1982; Ehleringer, 1993; Hall et al., 1994; Rebetzke et al., 2008; Des Marais et al., 2014; Raman et al., 2019), with some exceptions where nil or weak associations were observed (Hammer et al., 1997; Monneveux et al., 2007; Devi et al., 2011; Raman et al., 2020b).

Canola, being the second most important oilseed crop grown worldwide with a global production of 75 million tons (FAO STAT, http://www.fao.org/), is often cultivated in arid and semi-arid regions and faces periodic drought. Despite its economic significance to the oilseed industry as well as being an essential rotational crop in agricultural production systems, little research has been conducted on traits contributing to its drought tolerance (McVetty et al., 1989; Knight et al., 1994; Matus et al., 1995; Fletcher et al., 2015; Fletcher et al., 2016; Pater et al., 2017; Hossain et al., 2020; Raman et al., 2020a; Raman et al., 2020b). More recently, it was shown that two canola inbred lines, BC1329 and BC9102 vary by ∼ 2%_o_ in their Δ^13^C signatures (Hossain et al., 2020). However, the genetic basis of variation in Δ^13^C and other integrated WUE traits such as plant biomass, flowering time and seed yield was not deciphered. Thus, a comprehensive understanding of the genetic and physiological bases underlying WUE is central to developing strategies for resilience to water deficit conditions.

Herein, through comprehensive analyses based on extensive phenotypic and physiological measurements, genetic and genomic studies, we demonstrate that multiple genetic and environmental determinants underlie plasticity in multi-dimensional drought avoidance traits such as Δ^13^C, early vigour, plant height and seed yield, and drought escape traits such as flowering time in canola. We also show that one of the QTL for multi-traits; Δ^13^C, days to flower, plant height and seed yield on chromosome A09 is subjected to homoeologous recombination.

## Materials and methods

### Plant materials

In total, 223 doubled haploid (DH) lines derived from the F_1_ cross between advanced breeding lines ‘BC1329’ (maternal parent) and ‘BC9102’ (paternal parent) were utilised for different genetic analysis experiments. An F_2_ population comprising 744 lines derived from a single F_1_ plant from BC1329/BC9102 was employed for fine mapping/verification of QTL associated with Δ^13^C.

### Phenotypic evaluation for WUE traits

Four experiments were conducted in order to (i) determine the genomic regions that influence the expression of the traits associated with WUE (Experiments 1-3) and (ii) determine the relationship between Δ^13^C, *i*WUE and integrated WUE related traits under wet and dry conditions (Experiment 4). Experiments 1 and2 were performed under natural field conditions to measure WUE at the plot level; Experiment 3 is a pot experiment for single plant level WUE measurements and Experiment 4 is a rain-out shelter experiment with wet and dry irrigation regimes for measuring WUE at the single leaf level. Details of the experimental designs are presented in Table S4. Monthly weather statistics for average atmospheric temperatures and rainfall are also presented (Fig. S1).

### Phenotypic trait measurements

Several agronomic, gas exchange and other physiological traits were measured for genetic analysis. A summary of the experiments in terms of their aim, trial layout, genetic material evaluated, and the traits measured are presented (Table S1, Fig. S2). Details of trait measurements are given in our recent study (Raman et al., 2020b) and summarised in Table S2. A brief description of the traits measured is given below.

#### Plant development and agronomic traits

Δ^13^C, flowering time, plant height and seed yield were measured for Experiments 1-4 and normalised difference in the vegetative index (NDVI) was measured only for Experiment 2. Δ^13^C was determined from multi-phase experiments with appropriate experimental designs (Smith et al., 2006)) to account for the variations attributed to field/pot and laboratory conditions. The δ^13^C composition was determined using Vienna Pee Dee Belemnite (VPDB) as the ultimate reference. Δ^13^C was calculated from the δ^13^C values assuming the isotopic composition of CO_2_ in the air to be -7.8‰ on the VPDB scale, as described previously (Farquhar & Richards, 1984). Fresh and dry weights of the leaf and leaf thickness were also measured from F_2_ plants and row plots under wet conditions in Experiment 4.

#### Physiological traits

The gas exchange measurements were taken at the single leaf level for the plots under wet conditions in Experiment 4, as this relationship varies under different water-deficit levels. We determined *i*WUE by measuring light-saturated assimilation rate (*A*) and stomatal conductance to the diffusion of water vapour (*g*_*sw*_). The 5^th^ fully expanded leaf of each of the 72 lines of BC1329/BC9102 DH population (06-5101DH), including parental lines, was tagged and utilised for gas exchange measurements.

### Light microscopy

A leaf disc (9.08 cm^2^ size) was taken from each of two replicate canola lines from Experiment 4 (wet block), fixed and stored in 70% ethanol as detailed (Table S4). Leaf sections were stained using a method modified from Rae *et al*. (2020) and were imaged using 488 nm excitation and 500-560 nm emission on a Leica SP8 confocal microscope.

### Genotyping and linkage map construction

Genotyping of DH lines was carried-out using the genotyping-by-sequencing (GBS) based DArTseq approach (Raman et al., 2014). Sequence polymorphisms were used for linkage map construction following the method detailed in Raman *et al* (2016). The markers that showed complete segregation between each other were ‘binned’ into a unique locus and the resulting ‘bin’ map was used to identify trait-marker associations. To obtain the physical position of markers, DArTseq sequences were aligned with the Darmor-*bzh* reference assembly version 4.1 using the default parameter settings with the Bowtie program.

### Statistical methods

Commensurate with the aims of the experiments and the structure of the data sets, for Experiments 1-3 whole genome, single-step quantitative trait loci (QTL) analyses were performed on each trait using an extension of the approach developed by Verbyla and Cullis (2012) within a multi-environment trial (MET) analysis framework using factor analytic linear mixed models (FA-LMM) (Smith et al., 2015). Whereas, each trait measured on Experiment 4 is analysed individually using appropriate linear mixed models (LMM). A detailed description of the methods is presented (Table S4).

All analyses were performed in *ASReml-R* (Butler et al., 2018), which provides residual maximum likelihood (REML) estimates of variance parameters, empirical best linear unbiased predictions (EBLUPs) of random effects and empirical best linear unbiased estimates (EBLUEs) of fixed effects. The extent of genetic control of traits was investigated by calculating line mean *H*^2^ (broad-sense heritability) as the mean of the squared accuracy of the predicted DH line effects as described previously (Cullis et al., 2006) and found to be dependent on the environment. The across environment summary measure of Overall performance (OP) proposed by Smith and Cullis (2018) was used to identify lines of interest. We examined the relationships of Δ^13^C with agronomic traits (seed yield, days to flowering, plant height and NDVI) using pair-wise correlations of Overall performance estimates from the MET analysis of each trait or the EBLUPs from the LMM analysis of each trait.

### Identification of candidate genes for WUE

*Arabidopsis thaliana* genes which had been annotated with various WUE-related terms were retrieved from the TAIR 10 database (https://www.arabidopsis.org/). These genes were then used to identify putative homologues in canola.

### Resequencing and structural variation analysis of parental lines

Libraries from high-quality genomic DNA from both parental lines, BC1329 and BC9102, were constructed using the Illumina TruSeq DNA preparation kit, following the manufacturer’s instructions (Illumina). Whole-genome resequencing (2 × 150 bp) was performed at the Novogene facility (Novogene Co., Ltd, Hong Kong) using the Illumina HiSeq 2000 sequencing platform. The coverage of the parental lines ranged from 77.6× (BC1329, 102.6 Gb) to 83.8× (BC 9102, 112.4 Gb). Read mapping to the *‘*Darmor-*bzh’* reference assembly (version 4.1, http://www.genoscope.cns.fr/brassicanapus/data/), SNP and InDel (< 50-bp) calling, structural variation (SV, ≥ 50-bp) detection and identification of HE event (≥ 10-kb windows) was performed as described in Raman *et al*. (2021).

### Development of sequence-capture based DArTAg markers

We processed sequence data for target QTL regions on A09 and C09 chromosomes (Table S12) and selected 154 SNPs for DArTag oligo-synthesis. Oligos were synthesised by IDT (Ultramer DNA Oligos, http://idtdna.com) at 200 pmol scale, pooled in the equimolar amount into a single assay and used for processing 8 plates of DNA with the F_2_ population and a control canola sample using a proprietary DArTag assay (Targeted Genotyping - Diversity Arrays Technology) using 384 plate format. For each plate, a sample of the pooled product was also run on agarose gel and compared against positive control before proceeding with the sequencing process. The libraries were sequenced on Illumina Hiseq2500 with an average volume of sequencing per sample at 43,225 sequencing reads (median at 46,389) and average read depth per assay at 280. Marker data were extracted using DArT PL’s proprietary algorithm deployed a plugin in KDCompute application framework (https://www.kddart.org/kdcompute.html).

### RNA sequencing and differential gene expression analysis

Parental lines, BC1329 and BC9102 of DH population were grown in three replicates under both wet (100% field capacity) and dry (50% field capacity) treatments in a glasshouse (Table S4). The clean sequence reads (100 bp single-end reads) for 12 samples that had per base sequence quality with >96% bases above Q30 were aligned against the *B. napus* reference Darmor-*bzh* (Version 4.1), using STAR aligner (v2.5.3a) (https://github.com/alexdobin/STAR/blob/master/doc/STARmanual.pdf). The raw counts of reads mapping to each known gene was used to perform differential expression analysis using edgeR (version 3.30.3) (https://bioconductor.org/packages/release/bioc/html/edgeR.html) using R version 4.0.3. A generalised linear model approach was then used to quantify the differential expression between the groups. The differentially expressed genes (DEGs) were obtained using a false discovery rate (FDR < 0.05). Heatmaps showing the expression pattern of genes in A09 and C09 QTL regions were produced using the ComplexHeatmap R package (Gu et al., 2016).

## Results

### Substantial genetic variation in Δ^13^C and other WUE traits

We observed high levels of genetic variation in Δ^13^C and other WUE related traits in the DH mapping population. The significant source of genetic variation was from the additive component (genetic markers), which ranged from 21.5% for NDVI to 79.1% for days to flower (Table S5, Additive M1, %). Broad sense heritability estimates for Δ^13^C and other integrated WUE related traits (plant height, NDVI, flowering time and seed yield) were variable, ranging from low (56%) to high (98%), depending on the nature of trait and growing environment (Table S6). Estimated additive and total (additive plus non-additive) genetic correlations between environments revealed that there are strong correlations between environments for both additive and total genetic variance with values greater than 0.89 and 0.83, respectively, for all traits (Table S7). Overall performance estimates for Δ^13^C ranged from 18.73 to 21.25‰ and displayed transgressive segregation among DH lines across phenotypic environments (Fig. 1a, Table S8). Up to 2.52‰ variation in Δ^13^C was observed among DH lines that equates to a 5-fold increase compared with the parental lines.

**Fig. 1:**
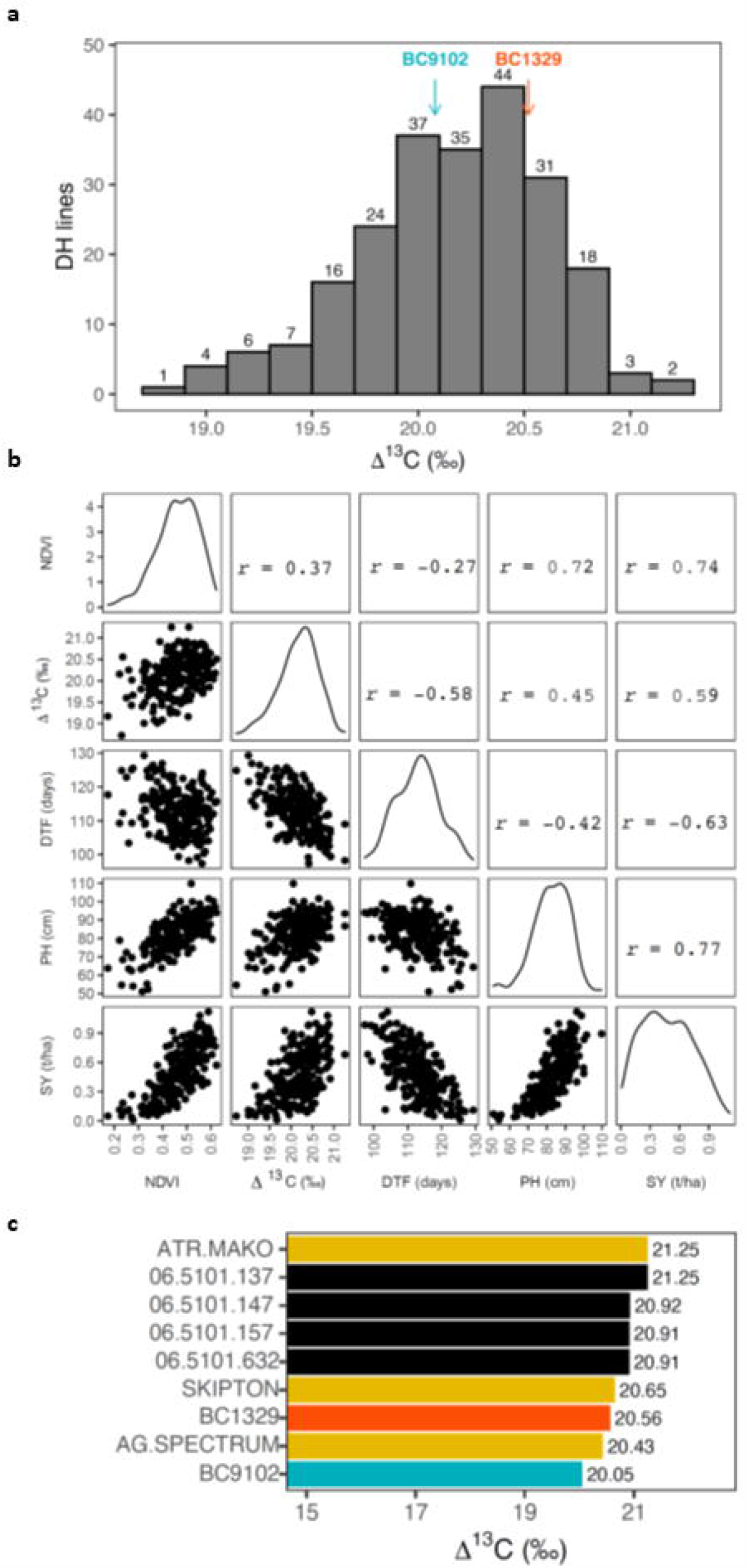
Genetic variation in WUE traits and their relationships among doubled haploid lines derived from the cross, BC1329/BC9102. **a:** Frequency distribution of the Overall performance estimates for Δ^13^C. Estimates for the parental lines are shown with arrows; **b:** Pair-wise correlations of the Overall performance estimates between Δ^13^C (‰) and other WUE related traits; **c:** Top four DH lines that showed the highest Δ^13^C based on Overall performance estimates across environments in relation to control commercial varieties of canola and the parental lines are shown.

### Relationships between WUE traits at plot level

To determine the relationships between Δ^13^C and other WUE related traits, pair-wise correlations were obtained using the genotype Overall Performance estimates across environments (Fig. 1b). The Δ^13^C showed a negative correlation with days to flower (*r* = - 0.58), while positive correlations were observed with NDVI, a proxy for plant vigour (*r* = 0.37), plant height (*r* = 0.45) and seed yield (*r* = 0.59). Flowering time showed a negative correlation with seed yield (*r* = - 0.63). The promising DH lines that had high WUE of yield (high Δ^13^C) for use in canola breeding programs based on the Overall performance estimates are presented in Fig. 1c. DH line 06-5101-137 had the maximum Δ^13^C (21.25‰) among the DH progenies.

### Relationships between physiological WUE (single leaf level) and integrated WUE (whole plant level)

Significant variation for both *A* and *g*_*sw*_ was observed, although *H*^*2*^ estimate of *i*WUE was low (Table S9). This may have occurred due to variable VPD across gas exchange measurements during the experiment, highlighting the plasticity of *i*WUE and Δ^13^C as traits. Genotype EBLUPs for *A* and *g*_*sw*_ ranged from 4.97 to 17.15, and 0.11 to 0.38, respectively (Table S9). Pairwise correlations revealed that both *A* and *g*_*sw*_ are dependent on each other with a correlation of 0.56 (Fig. 2a). We observed a negative correlation between Δ^13^C and *iWUE* (*r* = - 0.16), indicating that DH lines with low Δ^13^C have higher *iWUE*, consistent with the findings made earlier (Farquhar & Richards, 1984). There was a more negative correlation between *iWUE* and *g*_*sw*_ (*r* = - 0.46) in comparison to *A* (*r = -*0.27), suggesting that *g*_*sw*_ is the predominant driver for variation in *i*WUE parameters.

**Fig. 2:**
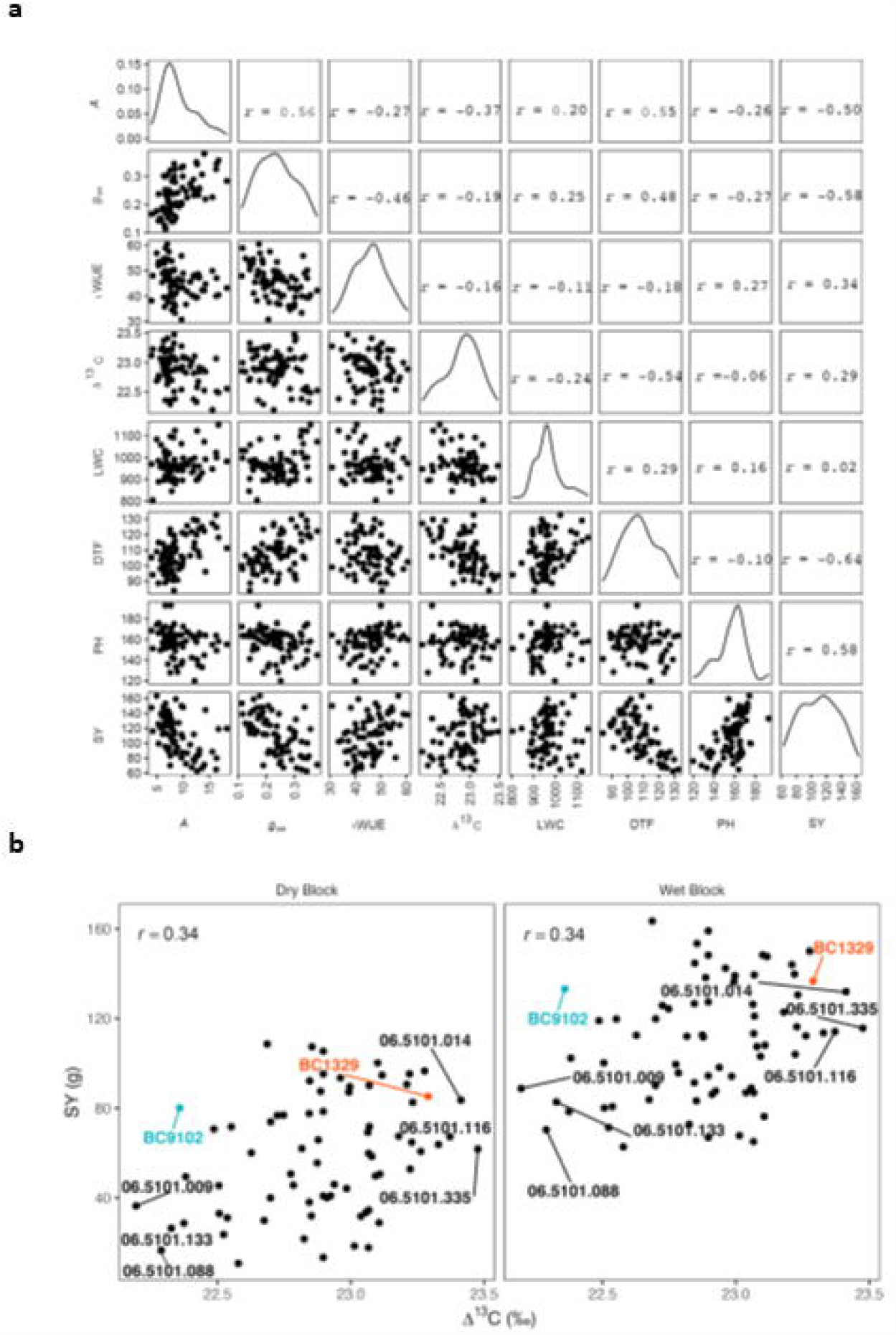
Relationships between Δ^13^C, gas exchange measurements (CO_2_ assimilation (A), stomatal conductance (g_sw_), and intrinsic water use efficiency (*i*WUE), plant developmental and agronomic traits (LWC: leaf water content; DTF: Days to flower; PH: Plant height and SY: Seed yield) of selected 70 DH lines of the BC1329/BC9102 population, representing extremes (High and low values) in Δ^13^C and their parents. **a:** Pair-wise correlations of the genotype EBLUPs are plotted. DH lines were grown under rain-out shelter with wet and dry conditions. **b**: Relationships between Δ^13^C and seed yield for wet and dry blocks. Genotype EBLUPs for Δ^13^C and seed yield are plotted. Parental lines and the DH lines with high and low Δ^13^C are labelled.

This study showed that Δ^13^C correlates negatively with *i*WUE but it (Δ^13^C) correlates positively with seed yield (Fig. 2a). Under well-watered conditions, there were negative correlations between Δ^13^C and days to flower, *A* and *iWUE*. We further investigated relationships between leaf water content (LWC) at a single leaf level and WUE traits at the whole plant level and found that LWC show a negative relationship with Δ^13^C, but it did not show any relationship with seed yield (Fig. 2a). Further, the estimated genetic correlations between wet and dry blocks for seed yield (Fig. 2b) and plant height, the only two traits measured after imposing water stress at the first flowering stage, were very high (0.93 for both traits). This suggests that genotype by irrigation block interaction is small. High Δ^13^C lines revealed higher yield across irrigation blocks compared to low Δ^13^C lines.

### Genetic basis underlying Δ^13^C and WUE related traits

We constructed a linkage map that includes 8,985 DArTseq markers onto 24 linkage groups (LGs), representing all the 19 chromosomes of *B. napus* (Table S10). To reduce computation time for genetic analysis, we produce a ‘bin’ map of 1793 markers that spanned a total of 1965.29 cM, with an average interval of 1.10 cM between adjacent loci.

Multi-environment QTL analysis identified a total of 29 QTL (15 QTL for main-effects and 14 for QTL (Q) × Environment (E) interactions) for variation in Δ^13^C and other WUE related traits (Table 1, Table S11). For leaf Δ^13^C, three QTL main effects that showed statistically significant (LOD ≥ 3) associations were identified on chromosomes A08, A09 and C09, while one ‘suggestive’ QTL (LOD >2.5 but less than 3) was located on chromosome A07 (Table 1, Fig. 3a). We identified QTL for phenotypic plasticity in different traits between three growing environments (Q × E effects) on A02, A05, A08, A09, A10, C02, C03, C06, C07 and C09 chromosomes (Table S11). For Δ^13^C plasticity, two QTL were identified on chromosomes A02 and C06, although the size of allelic effects were environment-dependent (Table S11). Collectively, QTL explained 38% of genotypic variation in Δ^13^C (Table S5, VAF_m_).

**Table 1:**
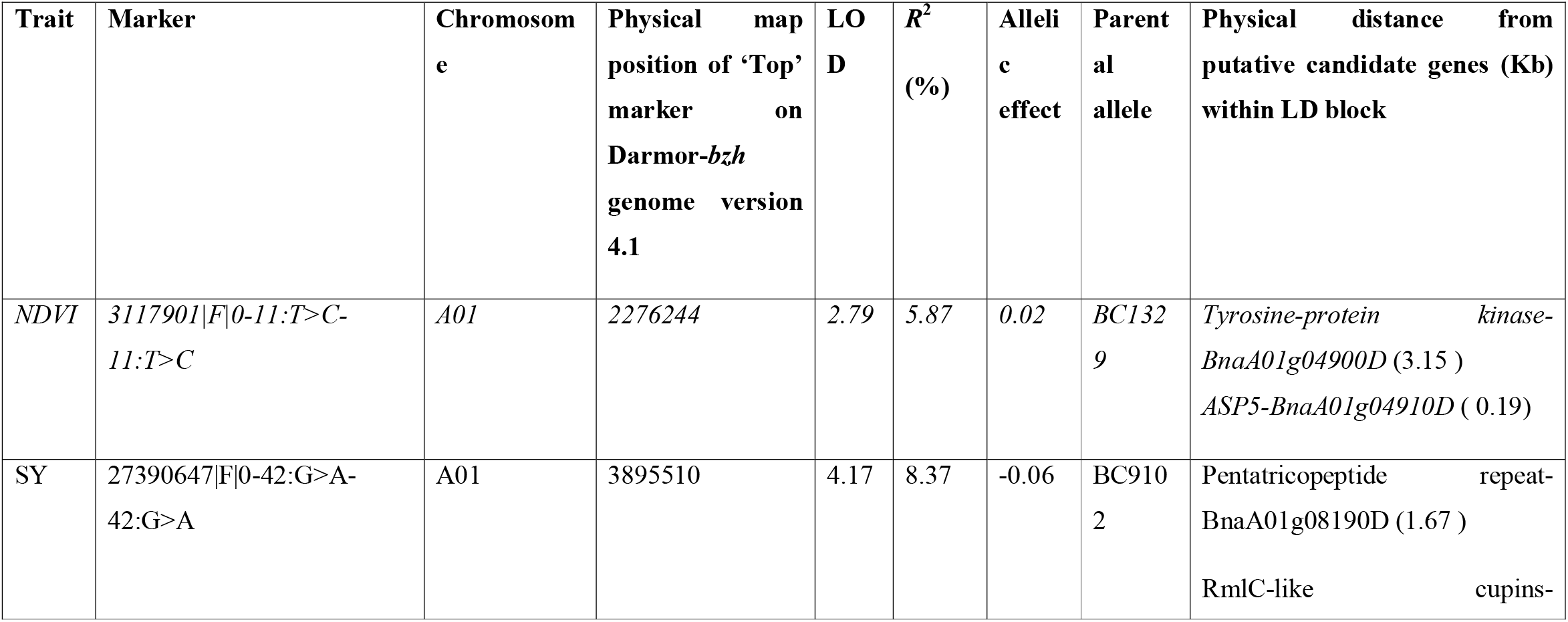
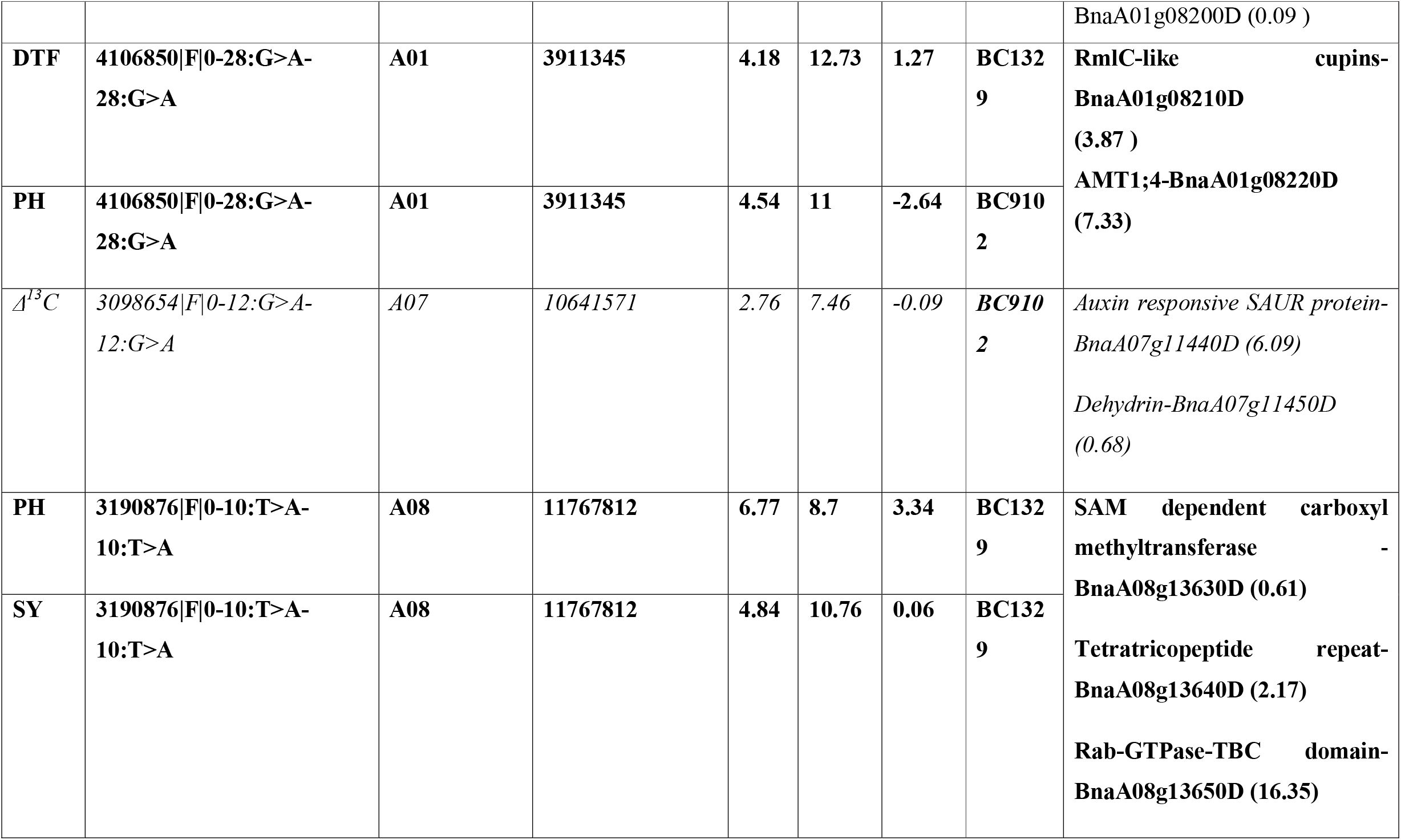
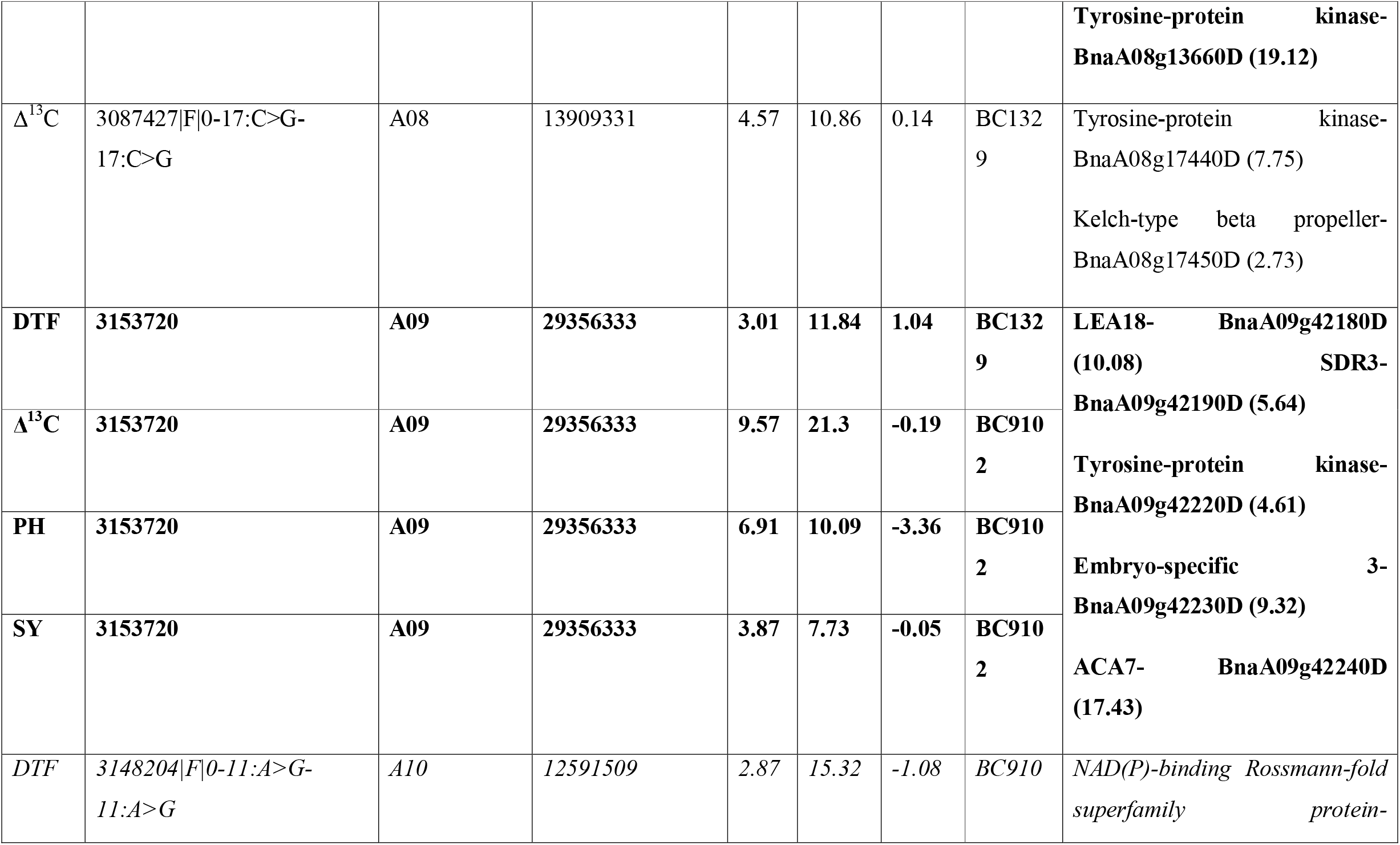
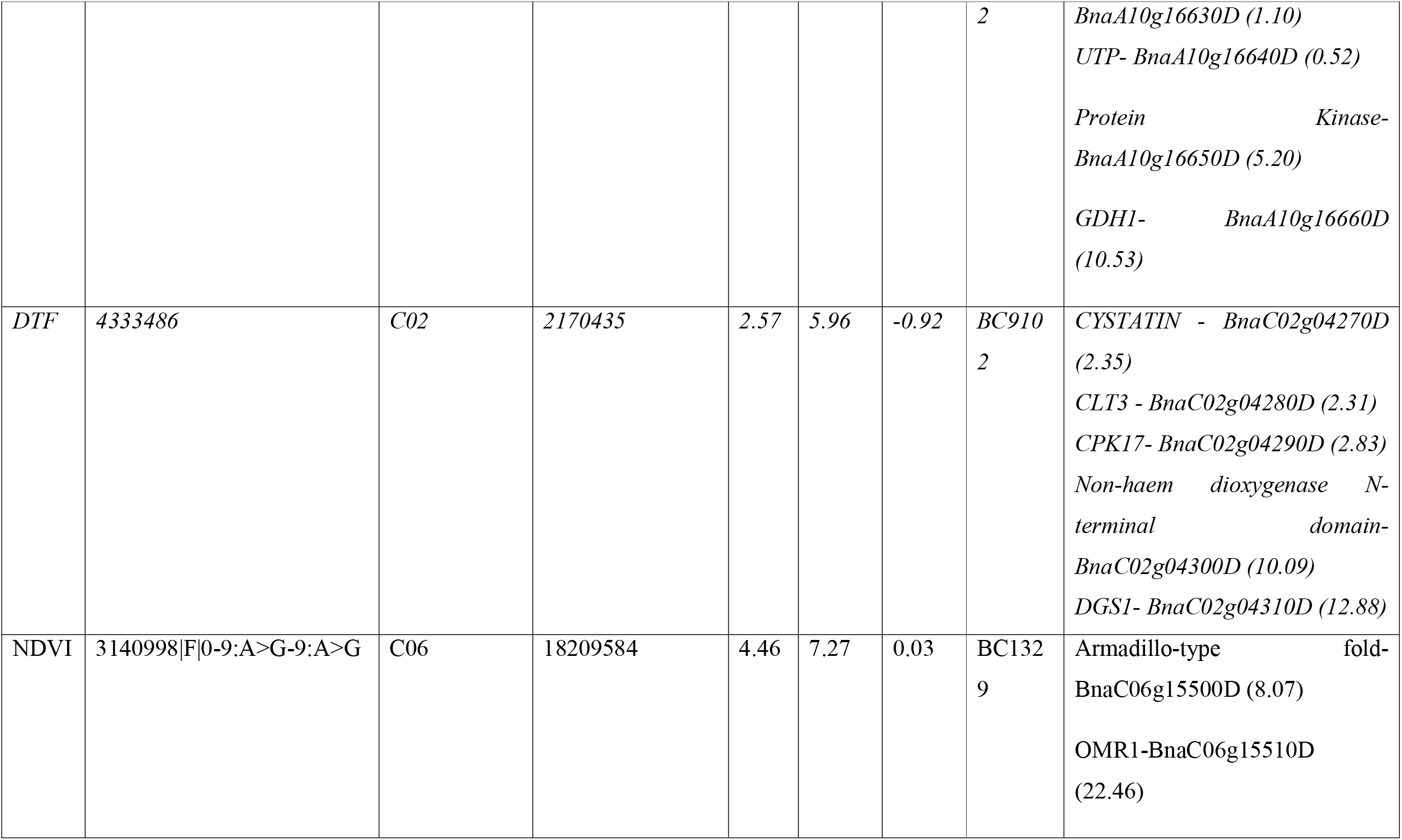
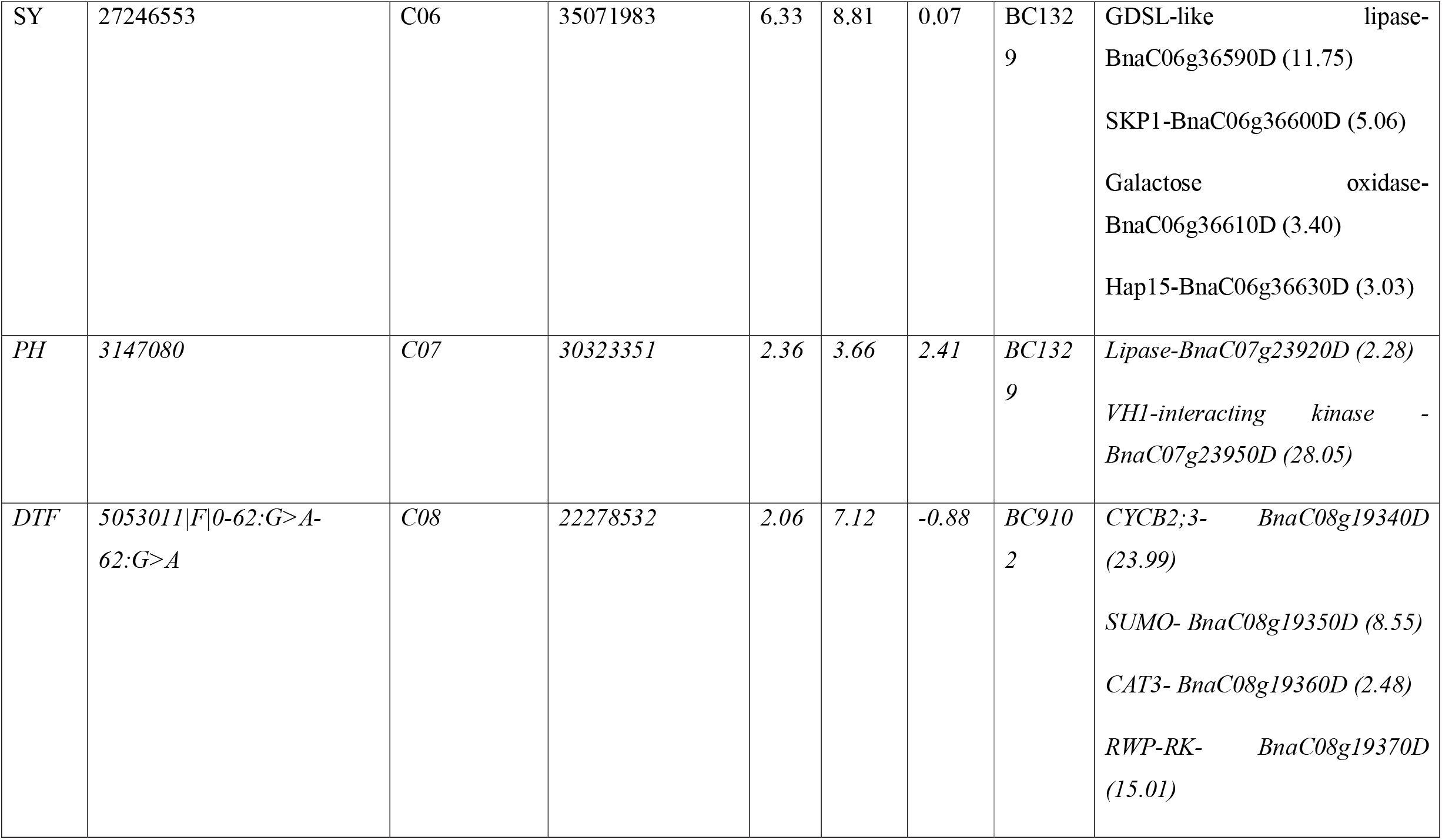
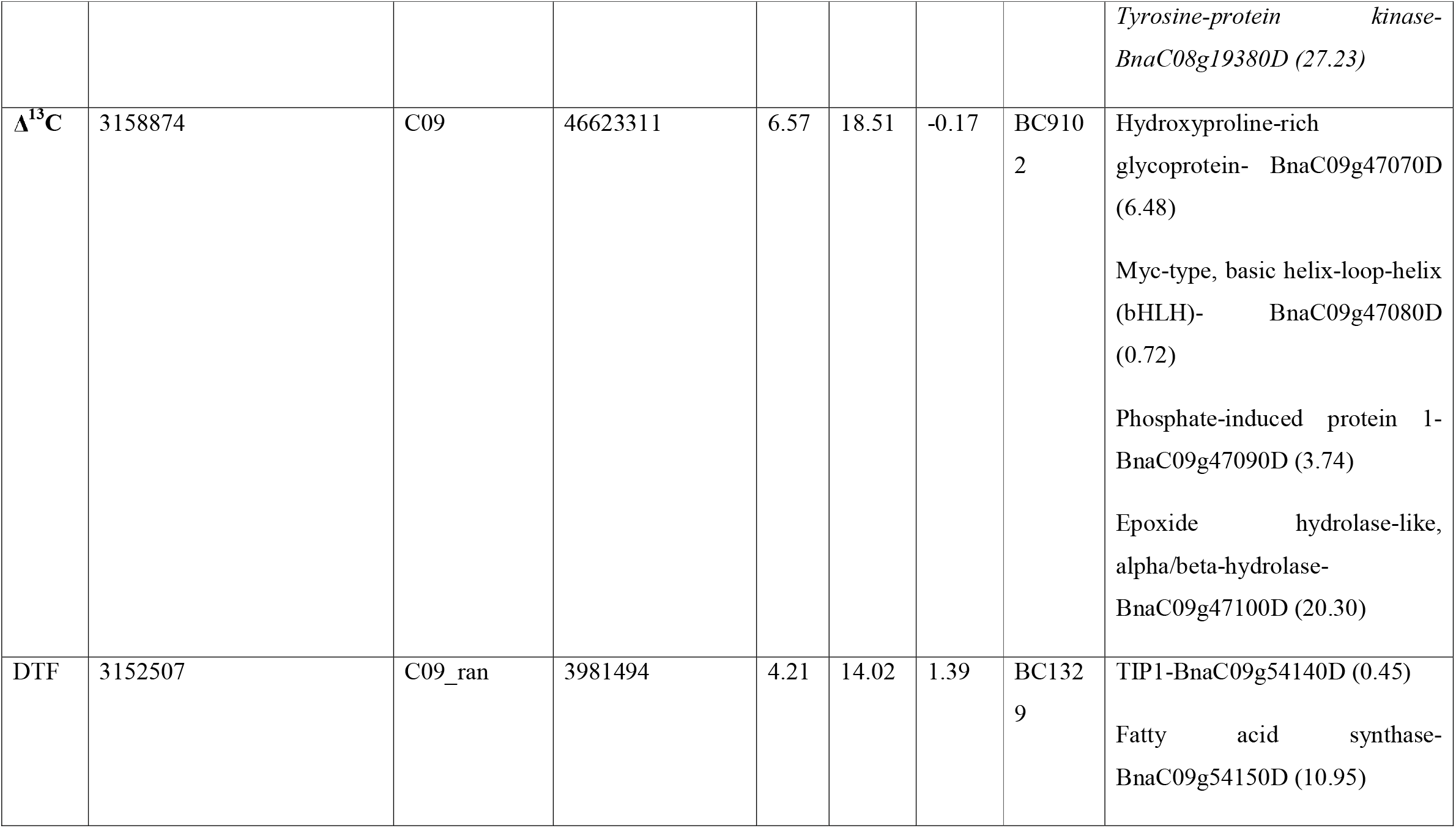
Quantitative trait loci (main effects) for carbon isotope discrimination (Δ^13^C) and agronomic traits (DTF: Days to flower; NDVI: Normalised difference vegetative difference; PH: Plant height; SY: Seed yield) evaluated in doubled haploid lines from BC1329/BC9102, across three environments. LOD scores, allelic effect, parental allele and percentage of genetic variance explained (*R*^2^) were also provided. QTL x Environment interactions for each environment are presented in supplementary Table S11. Putative candidate genes underlying QTL x Environment interactions are given in Table S13. Suggestive QTL having LOD ≤ 3 are in italics whereas consistent markers that were associated with multiple traits are in bold font.

**Fig. 3:**
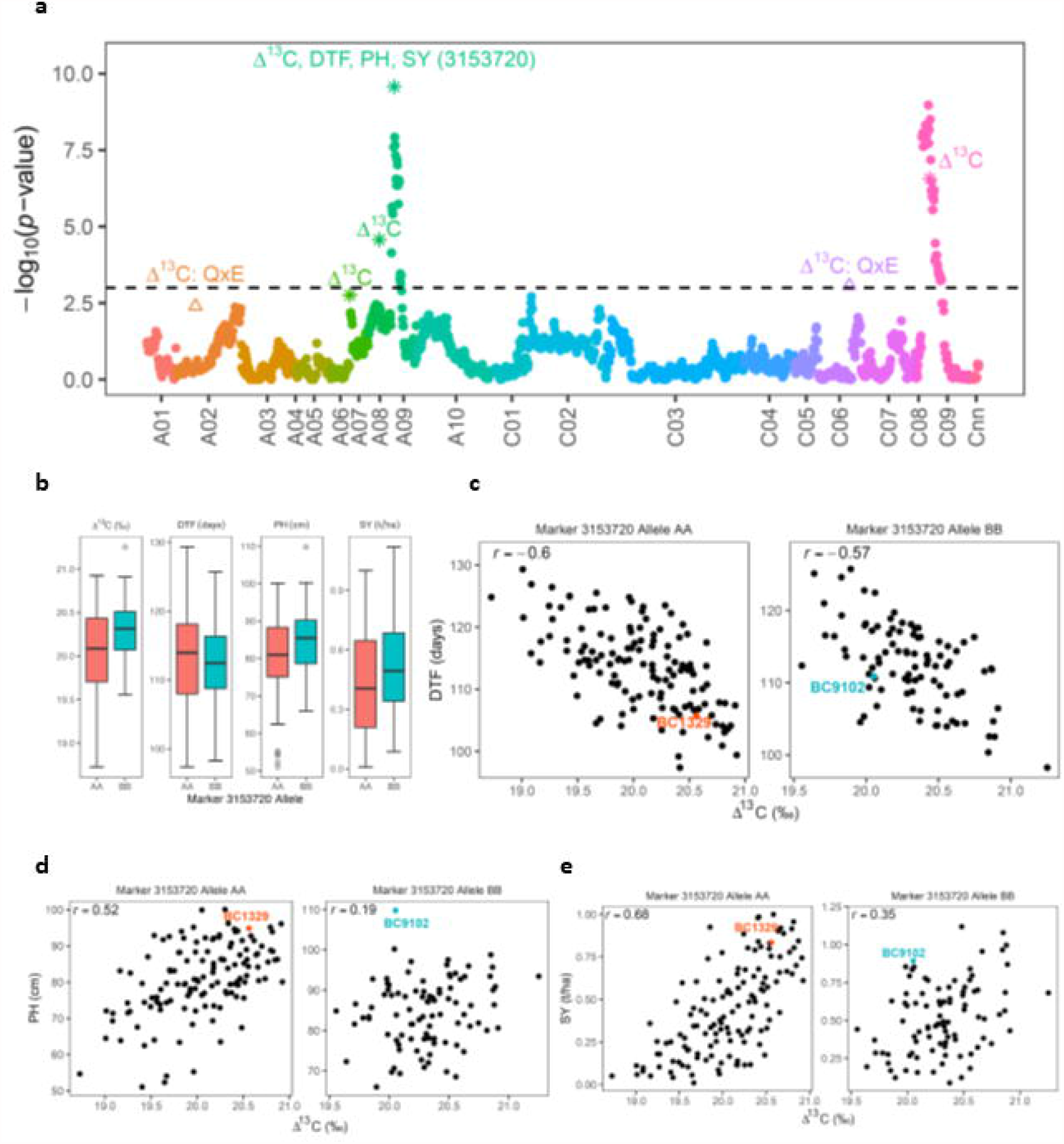
Distribution and relationships between Overall performance estimates of Δ^13^C, days to flower (DTF), plant height (PH) and seed yield (SY) and DArTseq marker alleles for the QTL (3153720) that colocalized in the same genomic region on chromosome A09. Manhattan plot showing LOD scores for associations between DArTseq markers and Δ^13^C (a). QTL main effects are labelled with the respective trait (for days to flower, plant height and seed yield only the 3153720 QTL is shown) and QTL x Environment interactions are labelled with the trait followed by ‘Q × E’ (only shown for Δ^13^C). LOD scores presented in the Manhattan plot are from the genome scan for the QTL main effects where the LOD scores of the significant QTL are replaced with the ones from the final model. The black dash line indicates the threshold value for significant SNPs at LOD ≥ 3. Box plots showing the distribution of the Overall performance estimates for Δ^13^C, days to flower, plant height and seed yield partitioned into allele combinations, ‘AA (BC1329)’ and ‘BB (BC9102)’, for the SNP marker 3153720 (**b**). Pair-wise correlations of Overall performance estimates between Δ^13^C vs days to flower (**c**), Δ^13^C vs plant height (**d**) and Δ^13^C vs seed yield (**e**) are partitioned into different allelic combinations.

### Comparative localisation of QTL

Three QTL for multi-traits on chromosomes A01, A08 and A09 were colocalised to the same genomic regions (Table 1). One QTL delimited with marker 3153720 for variation in Δ^13^C was colocated with days to flower, plant height and seed yield on chromosome A09 (Table 1, Fig. 3a). We further sought a correlation between allelic effects of markers and variation in Δ^13^C, days to flower, plant height and seed yield (Fig. 3b-e). Up to 68% of allelic effects were explained by the same marker allele (Fig. 3e), suggesting pleiotropic relationships between these traits and/or tight genetic linkage between them.

### Verification of QTL for Δ^13^C

We validated the genetic control, the linkage between DArTseq markers and Δ^13^C (in DH population) and focused on the identification of candidate gene(s) underlying the majority of genetic variation in Δ^13^C at QTL regions on chromosomes A09 and C09 (Table 1). The Δ^13^C values showed a wide range distribution among F_2_ lines (Fig. 4a). Unlike DH lines, Δ^13^C exhibited a positive correlation with flowering time and LWC, and a negative correlation with SLW (Fig. 4b-d). Our anatomical analysis of leaf discs that revealed both parental lines BC1329 and BC9102 differ in thickness and arrangement of palisade and spongy mesophyll cells: BC1329 (192 μm) had high porosity with large airspaces compared to BC9102 (184 μm, Fig. 4e-f), which may facilitate gas exchange, thus leading to efficient water use.

**Fig. 4:**
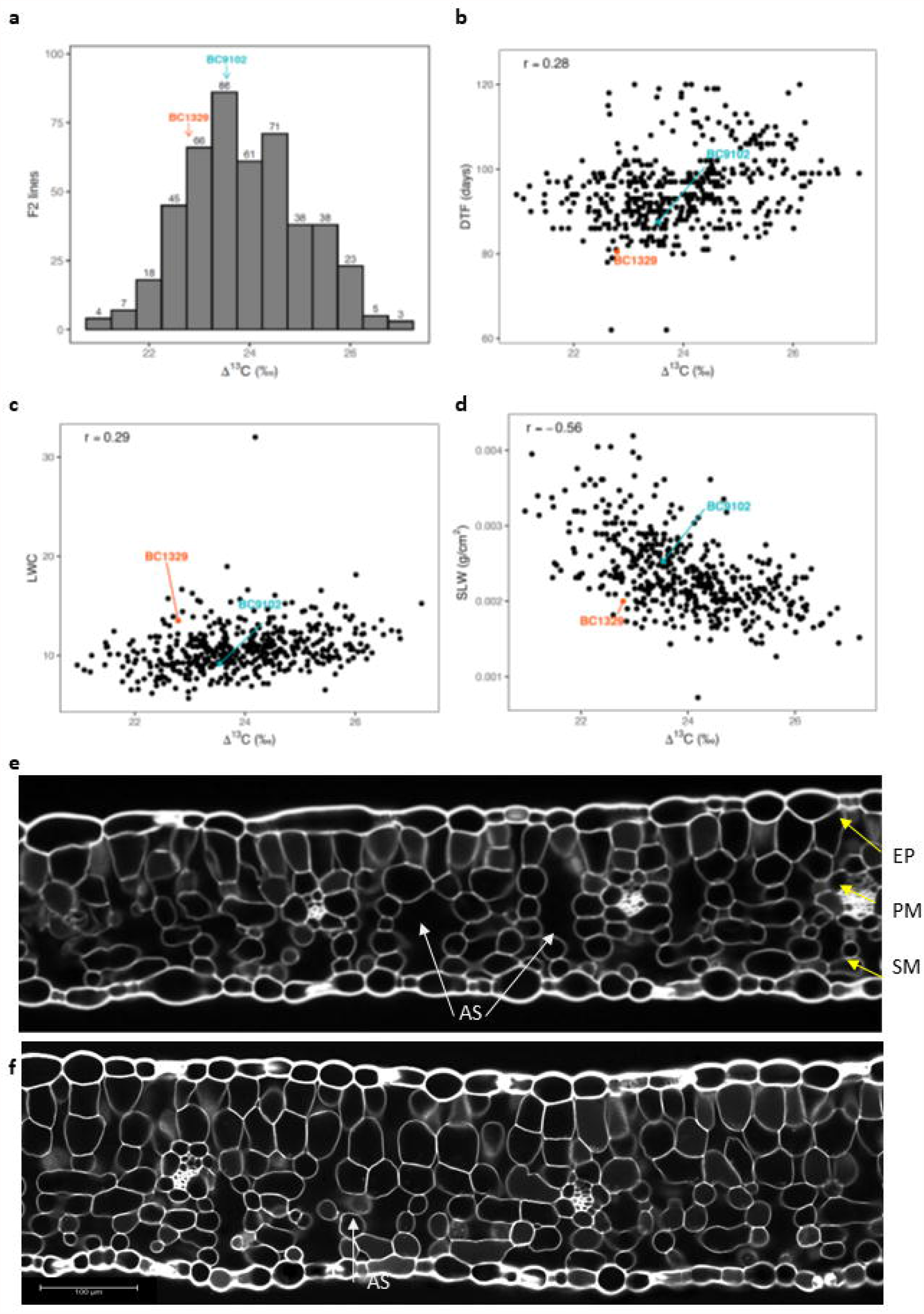
Distribution and relationships of the traits measured for an F_2_ validation population derived from the BC1329/BC9102, grown under non-stress conditions. The frequency distribution of Δ^13^C (‰) among 744 F_2_ lines (**a**). Pair-wise correlations between Δ^13^C and DTF (**b**), Δ^13^C and LWC (**c**) and Δ^13^C and SLW (**d**) are shown. Δ^13^C: Carbon isotope discrimination; DTF: Days to flower; LWC: Leaf water content; SLW: Specific leaf weight. Leaf sections showing differences in air spaces (AS, marked with arrow) between parental lines BC1329 (**e**) and BC9102 (**f**). EP: epidermis; PM: palisade mesophyll (comparatively regular elongated cells); SM: spongy mesophyll (irregular cells)

Genetic analysis revealed that several DArTag markers show significant segregation distortion (deviating from the normal segregation consistent with 1:2:1 ratio for codominance, or 3:1 ratio for dominance) on chromosomes A09 and C09 (Table S12), suggesting that the Δ^13^C region could be subjected to structural variation. Genome scan using linear marker regression revealed that DArTag markers positioned at 28,598,612 bp on chromosome A09, and 46318271 bp on C09 of the Darmor-*bzh* genome exhibit statistically significant association with Δ^13^C variation (Fig. S3).

### Physical mapping and candidate genes associated with WUE near Δ^13^C QTL

To identify potential candidate genes involved in the Δ^13^C variation, we interrogated genomic regions underlying the significantly associated markers in both the mapping (DH) and validation populations (F_2_). In the DH population, DArTseq 3153720 ‘bin’ marker revealed the complete linkage with another 12 markers, which were localised within 1.49 Mb region, spanning 28.35 Mb to 29.35 Mb (Table S10, Fig. S3). Annotation of genomic interval revealed that several genes including *ERECTA* (BnaA09g40540D), *PYL2* (BnaA09g40690D), *H*^*+*^*ATPase-5* (BnaA09g41340), *LEA18* (BnaA09g42180D) and *Protein Kinase* (BnaA09g42220D) on chromosome A09 and on its homoeologous chromosome C08, and *HAC11* (BnaC09g46960D), floral repressor *FLC (FLC*.*C09a*; *BnaC09g46500* and *FLC*.*C09b*; *BnaC09g46540D*), Myc-type BHLH (BnaC09g46950D, BnaC09g47080D on homoeologous group C09/A10 chromosomes are likely candidates to be involved in Δ^13^C variation (Table S13, Fig. S3). DArTag marker (physical position on the Darmor-*bzh* genome: 28,598,612 bp) on chromosome A09 was located within 93 kb of the *ERECTA* gene that controls transpiration efficiency in *A. thaliana* (Masle et al., 2005).

### Δ^13^C QTL region on chromosome A09 is subjected to homoeologous exchange (HE)

We observed significant segregation distortion among marker alleles on chromosomes A09 and C09 in both mapping (DH) and validation (F_2_) populations and inconsistency in collinearity across both genetic and physical maps (Table S12). To investigate whether QTL region on A09 is subjected to structural variation, we performed HE analysis utilising resequencing data of the parental lines. Sequence mapping revealed 26 genomic regions undergone HE events, varying from 90 kb to 870 kb, including the A09 multi-trait QTL region (29.3 to 29.5 Mb), BC9102 from C08 chromosome, as a result of homoeologous recombination (Fig. 5a, Table S14). However, in the maternal line BC1329, no such event was identified (Fig. 5b).

**Fig. 5:**
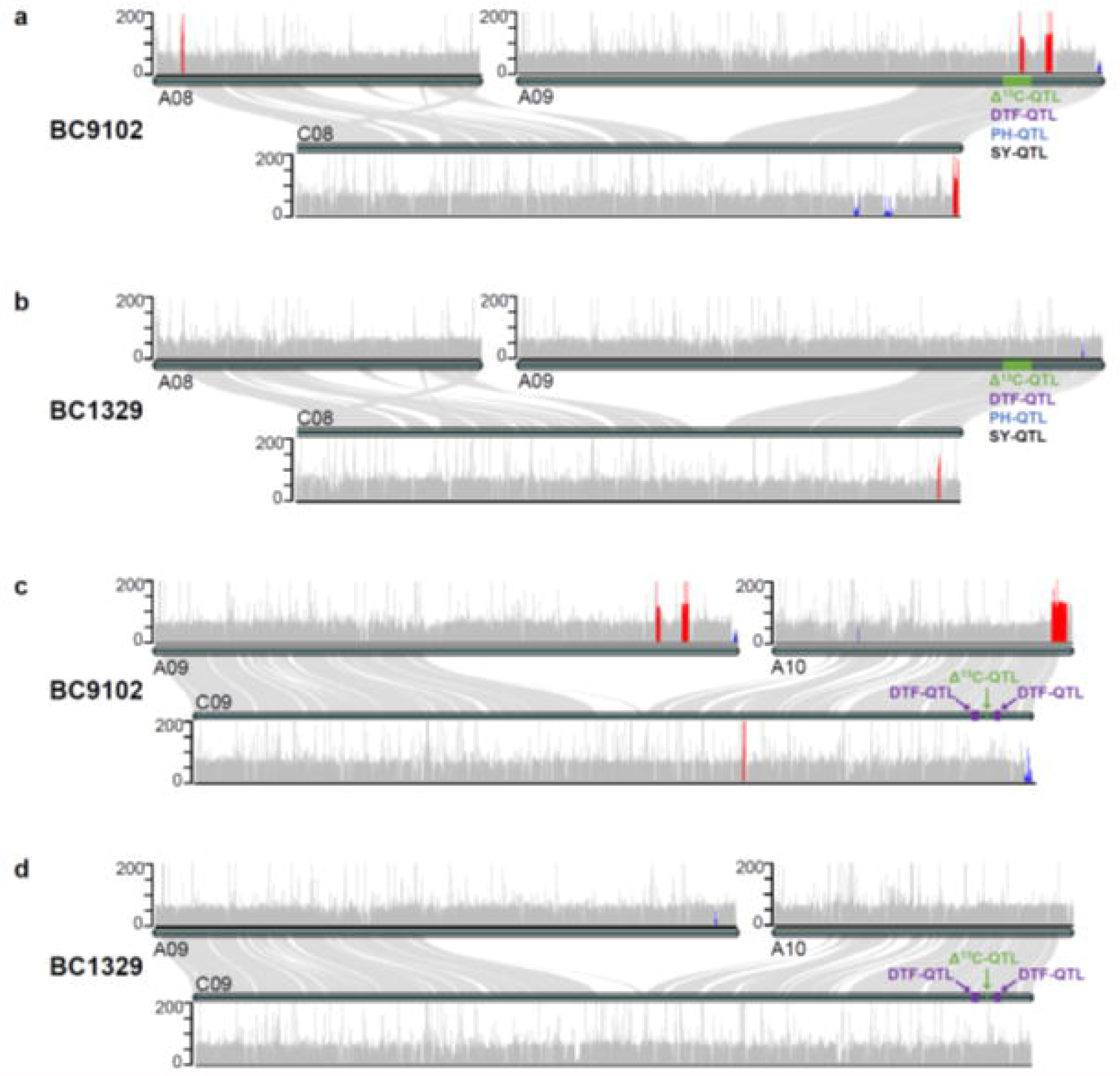
Homoeologous exchange (HE) events detected between parental lines of doubled haploid population derived from the BC1329/BC9102. Genomic sequences that undergone HE are shown in Table S14. Substituted and ‘translocated’ reads are highlighted in Blue and Red colour, respectively.

### Gene expression changes for Δ^13^C variation in the A09 and C09 QTL intervals between the parents

To investigate the expression of candidate genes that underlie the Δ^13^C variation on chromosomes A09 and C09, we examined the leaf tissue-specific transcriptome of the two parental lines: BC1329 and BC9102 under wet and dry conditions. We found that a total of 60 genes on A09 and 51 genes on C09 underlying Δ^13^C QTL regions were significanty differentially expressed between the two parental lines (Table S15). Of the DEGs, several of them such as Casein Kinase 2 α4 (BnaA09g42220D), Cation-transporting P-type ATPase (BnaA09g41340D, BnaA09g42040D), BEL1-like homeodomain protein 4 (BnaA09g41850D), Spermidine disinapoyl acyltransferase (BnaA09g41960D), Protein Kinase (BnaA09g42220D, BnaA09g41970D), HEC3 (BnaC09g46950D), and serine carboxypeptidase (BnaC09g47000D), are related with water use, water use efficiency and response to water stress (https://www.arabidopsis.org/). We also found that the expression levels of genes in BC9102 (with HE event) such as BnaA09g41850D, BnaA09g41970D (wall-associated receptor kinase-like 14), BnaA09g41990D (cyclin-dependent kinase inhibitor), BnaA09g42000D (nicotinate phosphoribosyltransferase 2), BnaA09g42030D (RNA recognition motif domain), and BnaA09g42040D were significantly higher (at least 2-fold) than those of BC1329 (without HE event) (Fig. 6, Table S15), suggesting that HE may be responsible for expression variation at the Δ^13^C-QTL region on A09.

**Fig. 6:**
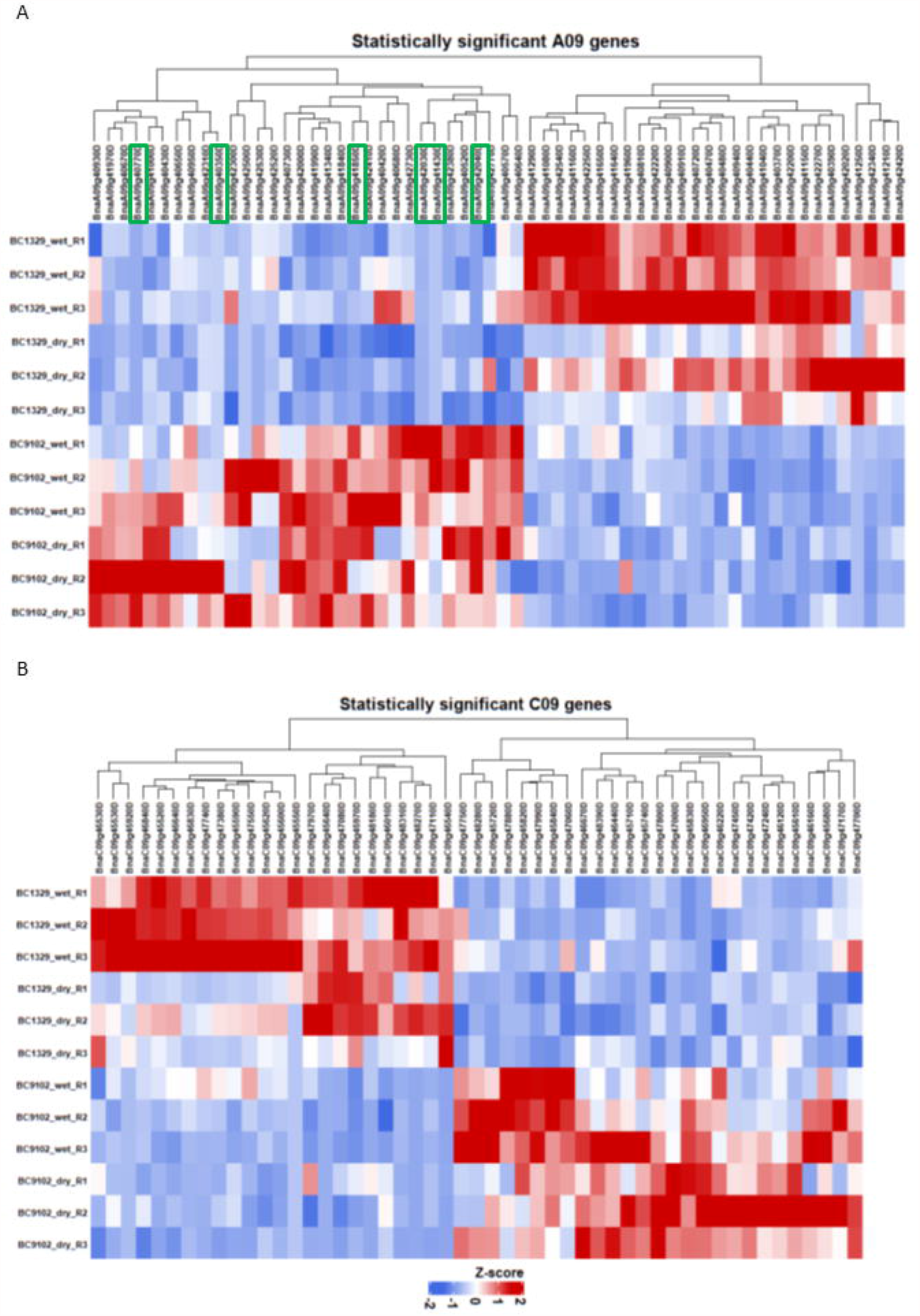
Expression profiles of differentially expressed genes (DEGs) in A09 (a) and C09 9b) QTL regions under water-deficit and water non-deficit conditions of the parental lines of the doubled haploid population derived from the BC1329/BC9102. The normalised read counts were plotted as a heatmap and genes were clustered according to the basis of their expression pattern. The genes in the heatmap were subjected to homoeologous exchange (HE) as well as the genes map within QTL region for Δ^13^C. DEGs that map within HE regions are highlighted in green boxes.

## DISCUSSION

### Canola reveals considerable variation for Δ^13^C

We found substantial genotypic variation in Δ^13^C, from 18.78 to 21.23‰ among DH, and 20.9 to 27.2‰ among F_2_ lines. An earlier study has shown that an increase of 0.5‰ in δ^13^C can lead to 25% more transpiration efficiency (TE = biomass gained/water transpired) in *Arabidopsis* (Juenger et al., 2005). Extrapolating this relationship, which is positive between δ^13^C and TE, and negative between Δ^13^C and TE, canola F_2_ lines with 6.3‰ higher Δ^13^C values than parental lines (22.8 to 23.5‰) should reduce WUE theoretically by 315%, which is impossible. It reflects the dependence of the sensitivity on the general level of Δ^13^C. For example, Masle et al. (2005) found that at the level they saw in *Arabidopsis*, an increase in Δ^13^C of 1‰ was associated with a 15% decrease in TE. Previous studies revealed that canola lines display a range of variation in Δ^13^C (18.7 to 23.7‰). Triazine tolerant (TT) accessions show higher Δ^13^C values compared to conventional open-pollinated varieties and hybrids (Matus et al., 1995; Pater et al., 2017; Hossain et al., 2020; Raman et al., 2020b). In this study, we utilised non-TT accessions for genetic analysis. Our research thus provides an additional genetic resource for understanding the genetic and physiological basis, as well as improving WUE in canola.

### Integrated WUE is partly driven by fitness traits

This study showed that DH lines that discriminate less between ^12^C and ^13^C as carbon source for photosynthesis (low Δ^13^C) show higher *i*WUE at the single leaf level (Fig. 2a). However, low Δ^13^ lines did not produce high yield (agronomic WUE; seed yield/unit of water used at the whole plot level) suggesting that selection for low *i*WUE at a single leaf level is useful for improving seed yield (*r* = 0.34, Fig. 2a), rather than using low Δ^13^C as a surrogate trait for predicting high seed yield in canola, consistent with our earlier findings (Raman et al., 2020b). This inconsistent relationship between Δ^13^C and seed yield could be due to genotypic variation in WUE being driven by variation in water use rather than by variation in assimilation per unit of water applied (Kobata et al., 1996; Blum, 2005; Sinclair, 2018). WUE, being a multi-dimensional trait can also be driven with other ‘fitness’ traits that reduce evapo-transpiration rate and crop water use. For example, high Δ^13^C lines with faster growth (NDVI, a proxy for plant vigour and plant height) could provide quicker canopy cover, which enables plants to reduce water loss from soil evaporation, thus increasing seed yield (*r* = 0.45 to 0.72, Fig. 1b). This is partly supported by in this study showing high correlation between plant fitness and seed yield and tight linkage of corresponding QTL (Fig. 1-3). In addition, Δ^13^C exhibited negative correlations with flowering time (*r*= - 0.58; DH population), and a positive correlation with NDVI, plant height and seed yield (Fig. 1b), suggesting that high Δ^13^C lines tend to ‘escape’ via accelerating growth and flowering - an evolutionary trait for adaptation to terminal drought stress. Our results showed that genotypes with low Δ^13^C had less canopy cover, late flowering and lower seed yield; these characteristics are typical for plants with drought avoidance strategy (TE). However, under terminal water-deficit situations, low Δ^13^C lines could yield poorly due to the shorter seed filling period, accompanied with high temperatures. It remains to establish how low Δ^13^C lines which require a longer season for seed filling, perform in climates that are not prone to environmental constraints (non-water deficit/heat stress).

### Genetic and environmental determinants affect phenotypic trait expression

We observed plasticity between Δ^13^C, and flowering time evaluated under field/pot and rain-out shelter (negative correlation, Fig. 1b, 2A) but a positive correlation under glasshouse conditions (Fig. 4b). This could be due to growing conditions (non-water stress condition, 100% field capacity) and nature of leaf tissue (discs without much vascular tissue) analysed for Δ^13^C.

Our comprehensive multi-environment QTL analysis showed that by using well-designed multiphase experiments (Table S3), and efficient statistical models (Table S4), both genetic and environmental determinants underpinning phenotypic variation can be deciphered for traits of interest (Table S11). For example, we identified QTL for the main effects (on A01, A07, A08 and A09) and Q x E interaction effects (on A02 and C06) that describe Δ^13^C plasticity across different environments (Table S11). Multi-environment based QTL analysis is a more powerful approach to dissect complex traits than the traditional QTL approaches (Zhang et al., 2010) but it was not used to uncover the genetic basis of WUE traits in canola previously. Consistent detection of Δ^13^C-QTL across three environments suggests that these loci contribute to the adaptive capacity of DH lines to water-deficit stress conditions and thus translating to economic seed yield (∼1 t/ha). Across field environments, DH lines were subjected to water deficit conditions, right from stem elongation to seed maturity (rainfall ranged from 225 to 235 mm over seven months of growing season, Fig. S2). Colocation of QTL for seed yield, Δ^13^C and plant height at the same genomic regions and stable allele (BC9102), contributing to trait variation that suggest multi-trait QTL on chromosome A09 are associated with effective water use. Early flowering showed a negative relationship with seed yield (Table 1), reiterating crosstalk between drought stress signalling and flowering time pathways (Des Marais et al., 2012).

It was interesting that none of the Δ^13^C QTL that we identified for main effect and Q x E interactions (Table S11) were detected in the Skipton/Ag-Spectrum population (Raman et al., 2020b). In an independent study, Mekonnen *et al*., (2020) identified three QTL for δ ^13^C on chromosomes A02, A09, and C08 in the North American *B. napus* mapping population. However, none of the QTL were consistently detected across environments. It is yet to establish whether the genomic region on chromosome A09 or its homoeologous counterpart C08 (QTL for root pulling force, plant height and δ ^13^C) is the same as found in our study, as the authors did not report the physical positions of QTL marker-intervals. In addition, there was a poor marker coverage on chromosome C08 in our genetic mapping population (13 markers, Table S10), which may have led to QTL (if any) being undetected in the unmapped regions, especially in HE region. These studies suggest that several genomic regions on A02, A03, A07, A09, C03, C06, C08, and C09 control variation in Δ^13^C, thus, genetic architecture of Δ^13^C is rather complex.

### *A priori* genes regulating WUE and efficient water use underlie QTL for Δ^13^C

Coarse and high-resolution mapping approaches utilised herein facilitated the validation of genomic regions for Δ^13^ variation and delimited candidate genes in canola, which are implicated in leaf-level WUE (Hersen et al., 2008; Cutler et al., 2010; Youn et al., 2016; Tao et al., 2018; Menéndez et al., 2019). For example, this study identified and validated a QTL that influences multiple traits; Δ^13^C, days to flower, plant height and seed yield on chromosome A09 that map within 92 kb of the *ERECTA* gene (Table S13). In different plant species, *ERECTA* and *ERECTA Like 1,2* genes encoding leucine-rich repeat protein kinases, regulate stomatal density and patterning, inflorescence architecture, ovule development, transpiration, and thermo-tolerance (Torii et al., 1996; Godiard et al., 2003; Shpak et al., 2003; Masle et al., 2005; Meng et al., 2012; Pillitteri & Torii, 2012; Bemis et al., 2013; Shen et al., 2015; Guo et al., 2020). *ERECTA* is also shown to control spikelet number-a component trait of grain yield via crosstalk between a Mitogen-activated protein kinase (MAPK) signalling pathway and cytokinin metabolism in rice. However, we did not find any difference in the level of expression of *ERECTA* between parental lines differing in Δ^13^ (unpublished data). We also localised several stress-responsive genes, including DEGs that may contribute to drought avoidance strategies via signal transduction pathways, encoding functional proteins (LEA18, RD20, glycine metabolism, CAT) and regulatory proteins, including transcription factors (bHLH, MYB, TINY2, ATHB6), protein kinases (Tyrosine protein kinase, Wall-associated receptor kinase-like 14, MAPK, SNF1-related protein kinase) and receptors (ABA receptor PYL12), phosphatases (PP2C), and calmodulins (CPK17) (Jonak et al., 2002; Des Marais et al., 2014; Jagodzik et al., 2018; Yong et al., 2019) within QTL intervals associated with Δ^13^C variation (Table 1, Table S13, S15). Plant expressing PYL12, and SRK2C genes are shown to improve the water use and drought tolerance (Yang et al., 2016) whereas ABC transporter (ABCG22) and ABA responsive kinase gene, MPK12 reduced the WUE (Des Marais et al., 2014). Our data hint that genes affecting stomatal characteristics (RD20, ERECTA), leaf thickness and water-deficit responsive genes described above likely underlie WUE and drought avoidance traits, while Q x E interactions are likely driven by environmental cues (PHYTOCHROME C was mapped with 6.2 kb from Δ^13^C-QTL on C06, Table S13).

Our results suggest that a QTL region underlying Δ^13^C, flowering time, plant height and seed yield on chromosome A09 may be subjected to HE. Homoeologous recombination is associated with presence-absence variation (Nicolas et al., 2007; Hurgobin et al., 2018). Recently, a major QTL for homoeologous recombination, *BnaPh1* was mapped on A09 (Higgins et al., 2021) and this was located within 5 Mbp of the QTL region that is associated with multiple traits. It is possible that the same genomic region may be involved in regulating WUE in diverse canola accessions and require further research.

In summary, this current study demonstrates that measures of *i*WUE, Δ^13^C and integrated WUE are complex and modulated by environmental and genetic determinants, including those subject to homoeologous exchange. Our findings on identification of useful variation in Δ^13^C (up to 6.3‰) and its underlying basis of variation in WUE traits, including their plasticity across environments, and identification of favourable alleles for increasing WUE would provide potential resources for developing new drought tolerant varieties for drier-environments to continue making genetic gains in the breeding programs.

## Sequence data availability

The raw sequence data reported in this paper has been deposited in the National Center for Biotechnology Information Sequence Read Archive (Accession no. PRJNA743730 for RNA-Seq data, PRJNA743989 for whole-genome resequencing data).

## Acknowledgements

This study was supported by the Australian Grains Research and Development Corporation and NSW Department of Primary Industries and partners (projects: DAN00117, and DAN00208. We thank Dr Simon Diffey (Apex Biometry) for multi-phase experimental designs for Δ^13^C and Dr. Alison Smith (UOW) for constructive discussions on the statistical methods.The authors are grateful to Mr. Warren Bartlett and Mr. Dean McCullum, for their assistance in sowing and management of field experiments; Hannah Roe and Wayne Pitt for grinding leaf samples for Δ^13^C analysis and Advanta for providing F_1_ cross.

## Author contributions

HR designed research; HR, RR, BM and YQ, performed field experiments; HSW analysed samples for carbon isotope discrimination; BC developed the statistical methods; RP, YZ, NS, HR, and SL analysed data; HR, NS, AZ and AK developed DArTags; BM, HR and GF performed/interpreted gas exchange measurements; RW investigated microscopic analysis; AE and DT provided seeds of F_1_; HR prepared manuscript with inputs from others. All authors read and approved this manuscript for publication.

