## Supplementary material for "Multi-environment QTL analysis delineates a major locus associated with homoeologous exchanges for water-use efficiency and seed yield in allopolyploid *Brassica napus*": Table S1

Table S1: Description of phenotyping experiments conducted to uncover genetic basis of carbon isotope discrimination (Δ^13^C) and water use efficiency related traits in canola (*Brassica napus* L).

| **Experiment** | **Research question** | **Material** | **Phenotyping environment** | **Blocks** | **Rows** | **Ranges** | **Plots** | **Traits measured** |
| --- | --- | --- | --- | --- | --- | --- | --- | --- |
| 1 | Genetic basis of CID variation | 223 DH lines from BC1329/BC9102, plus parental lines | Field (2017) | 2 | 45 | 10 | 450 | Δ^13^C, days to flower, plant height, seed yield |
| 2 |  | 223 DH lines from BC1329/BC9102, plus parental lines and commercial cultivars | Field (2018) | 2 | 76 | 6 | 456 | Δ^13^C, plant vigour (NDVI), days to flower, plant height, seed yield |
| 3 |  | 217 doubled haploid lines from BC1329/BC9102, plus parental lines | Pot (2017) | 4 | 73 | 12 | 876 | Δ^13^C, days to flower, plant height |
| 4 | Relationship between physiological (intrinsic water use efficiency, CID) and agronomic water use efficiency (canola productivity) related traits | Selected 70 DH lines representing extreme (Low and High CID values) plus parental lines under wet and dry conditions | Rain-out shelter (2019) | 2 | 48 | 9 | 432 | Δ^13^C, days to flower, plant height, seed yield, Photosynthesis (*A*), stomatal conductance (*g_sw_*), intrinsic water use efficiency (*i*WUE), Specific leaf weight (SLW), Leaf water content (LWC), Leaf thickness |
