## Supplementary material for "Multi-environment QTL analysis delineates a major locus associated with homoeologous exchanges for water-use efficiency and seed yield in allopolyploid *Brassica napus*": Table S2

**Table S2:** Trait measurement protocols for the plant development, agronomic and physiological traits measured in Experiments 1-4.

| **Trait** | **Measurement protocol** |
| --- | --- |
| *Plant development and agronomic traits* | |
| Δ^13^C | For Experiments 1-3, leaf samples were taken from 10 plants of the plots at the vegetative stage (BBCH 40, prior to flowering) and processed as described earlier (Raman *et al.*, 2020b). For Experiment 4, leaf samples were taken only from the wet block. The Δ^13^C was measured separately for each experiment and in different “runs”, with each run consisting of 3 carousals and 49 samples in each carousal (including five standards at position 1st/2nd, 25th, and 48th/49th of the carousal). At least twenty percent of field plot/pot samples were duplicated for the measurement of Δ^13^C in order to account for variation due to the laboratory process and field/pot variation. |
| Flowering time | Recorded as days to flower when 50% of plants in each plot/pot showed the first flower. |
| Plant height | At maturity, for Experiments 1-2 and 4, five plants were randomly selected from the middle of each plot from and were measured from the soil surface to the top of main branch, including siliques. For the Experiment 3, plant height was measured for all 5 plants in each pot. |
| Seed yield | For Experiments 1-2, seed yield was measured on plot basis. Plots were harvested with a small plot header and seed yield was expressed in ton/ha. For Experiment 4, seed yield was expressed in gm and the number of plants per row were counted. |
| NDVI | Measurements were taken periodically from each plot in Experiment 2 using the GreenSeeker® (NTech Industries Inc., Ukiah, CA, USA) following manufacturer’s instructions. |
| Leaf thickness | This was carried out using 9.02cm^2^ (area of leaf disc) cake cutters. Fresh and dry weights were measured. |
| Leaf water content (LWC) | Determined as the difference between fresh and dry weights, expressed as a percentage of dry weight. Leaves were dried at 80°C for 48 hr, until they reached constant mass. Measurements were made from the same samples that were used for leaf thickness. |
| Specific leaf weight (SLW) | Expressed as leaf dry weight/leaf area. Leaf area was determined from digital scans of flattened excised leaves using the ImageJ program using thresholding and the magic wand tool (http://www.imagej.nih.gov./ij/). |
| *Physiological traits* | |
| Light-saturated assimilation rate (Photosynthesis, *A*) | The 5^th^ fully expanded leaf of a randomly selected plant in each plot in the wet block of Experiment 4 was tagged and gas exchange measurements were obtained using the gas exchange cuvette (LI-6400XT, LICOR Inc., Lincoln, NE, USA) between 10h and 16h (AEST) following conditions described in Raman et al (2020). The same 5^th^ leaf that was used for gas exchange measurements was used for Δ^13^C measurements. |
| Stomatal conductance to the diffusion of water vapour (*g_sw_*) |  |
| *i*WUE (*A*/*g_sw_*) | Expressed as the ratio between *A* and *g_sw_*. |
