## Supplementary material for "Multi-environment QTL analysis delineates a major locus associated with homoeologous exchanges for water-use efficiency and seed yield in allopolyploid *Brassica napus*": Table S3

**Table S3:** Details of multi-phase (field/pot and lab) experiments carried-out to measure Δ^13^C in the doubled haploid population from BC1926/BC9102.

| **Experiment** | **Phenotypic environment** | **Field/Pot phase design** | | | | | | | **Lab phase design** | | | | | | | | | | | | | | |
| --- | --- | --- | --- | --- | --- | --- | --- | --- | --- | --- | --- | --- | --- | --- | --- | --- | --- | --- | --- | --- | --- | --- | --- |
|  |  | **Rows** | **Ranges** | **Plots/ Pots** | **Genotypes** | **Genotypes with *p* replicates** | | | **Samples for Δ^13^C** | | | | | | **Runs** | **Carousals per Run** | | | | | | | |
|  |  |  |  |  |  |  |  |  | **Rows** | **Ranges** | **Plots/ Pots** | **Field plots/Pots with *q* replicates** | | **Total** |  |  |  |  |  |  |  |  |  |
|  |  |  |  |  |  | ***p* = 2** | ***p* = 4** | ***p* = 6** |  |  |  | ***q* = 1** | ***q* = 2** |  |  | **1** | **2** | **3** | **4** | **5** | **6** | **7** | **8** |
| 1 | Field (2017) | 45 | 10 | 450 | 225 | 225 | 0 | 0 | 45 | 10 | 447* | 366 | 81 | 588 | 4 | 3 | 3 | 3 | 3 | - | - | - | - |
| 2 | Field (2018) | 76 | 6 | 456 | 228 | 228 | 0 | 0 | 76 | 4 | 304 | 256 | 48 | 392 | 3 | 3 | 3 | 2 | - | - | - | - | - |
| 3 | Pot (2017) | 73 | 12 | 876 | 219 | 0 | 219 | 0 | 73 | 12 | 862* | 756 | 106 | 1078 | 8 | 3 | 3 | 3 | 3 | 3 | 3 | 3 | 1 |
| 4 | Rain-out shelter (2019) | 48 | 9 | 432 | 72 | 0 | 0 | 72 | 24 | 4 | 96 | 60 | 36 | 146 | 1 | 3 | - | - | - | - | - | - | - |

*Information not available for 3 field plots in Experiment 1 and 12 pots in Experiments 3.

The Δ^13^C was measured separately for each experiment and in different “runs”, with each run consisting of 3 carousals and 49 samples in each carousal (including five standards at position 1st/2nd, 25th, and 48th/49th of the carousal). At least twenty percent of field plot/pot samples were duplicated for the measurement of Δ^13^C in order to account for variation due to the laboratory process and field/pot variation. Field plots/pots were resolvable to carousal, i.e. duplicated samples (which were from the same field plots/pots) were not allocated to the same carousal. The standards used in the lab phase were excluded from the data for analysis
