## Supplementary material for "Multi-environment QTL analysis delineates a major locus associated with homoeologous exchanges for water-use efficiency and seed yield in allopolyploid *Brassica napus*": Table S4

**Table S4. Supplementary materials**

**Experimental designs**

(i) **Experiments 1-2**

The DH lines (n = 223), along with its parental lines (BC1329 and BC9102) and three commercial check cultivars (Skipton, Ag-Spectrum and ATR-Mako) were evaluated in two field experiments; Experiment 1 and Experiment 2 during the 2017 and 2018 canola growing seasons, respectively (autumn, April-November/December). Experiment 1 was sown on 18^th^ May 2017 at the experimental station of Wagga Wagga Agricultural Institute (WWAG) (-35.0598782017231, 147.31173621965058) and Experiment 2 was sown on 14^th^ May 2018 (-35.04540035261563, 147.34480619057837). Both experiments were conducted in rectangular arrays of plots; Experiment 1 with 45 rows by 10 ranges and Experiment 2 with 76 rows by 6 ranges. The trial designs for both experiments were randomised complete block designs (RCBD) with 2 row-wise blocks for Experiment 1 and 2 range-wise blocks for Experiment 2. The lines (treatments) were allocated to plots within blocks where the blocks were resolvable with 1 replicate plot of each line occurring in each block. The plots were of the same width of 2 m in both experiments, consisting 8 rows, spaced 25 cm apart. Lengths of the plots were 10 m in 2017 and 6 m in 2018. The seeds were sown at a depth of approximately 2.5 cm with Impact-in furrow fertiliser to protect against blackleg disease, caused by the *L. maculans*. The seeding rate was 1,400 seeds/plot for Experiment 1 whereas in Experiment 2 it was adjusted to 840 seeds for comparison with Experiment 1. Pre-emergent herbicides were used to control weeds.

(ii) **Experiment 3**

Experiment 3 is conducted in 2017 using plastic pots (size 15 cm diameter) arranged in a rectangular array of 73 rows by 12 ranges under an area covered by glass on top (-35.04722451361815, 147.33105481807226). Five seeds for each of the 217 DH lines, along with parental lines: BC1329 and BC9102 were sown on 6^th^ June 2017 in the pots and raised to the full maturity stage (BBCH scale 95). The trial design was a randomized complete block design (RCBD) with 4 range-wise blocks. The lines (treatments) were allocated to plots within blocks where the blocks were resolvable with 1 replicate plot of each line occurring in each block. The plants were watered using poly-pipe drip irrigation system (water flow rate: 3.8L/h emitters) and supplemented with liquid in-line fertiliser “Campbells Diamond Blue” (Campbells Fertilisers Australasia) as required. Plants received water three times a week for 15 minutes.

(iii) **Experiment 4**

To determine the relationship between Δ^13^C, *i*WUE (*A*/*g_sw_*) and agronomic WUE related traits such as plant height and seed yield, extreme DH lines were selected from the tails of the population, comprising 35 top and 35 bottom lines on the basis of their seed yield across the field experiments. These 70 DH lines together with their two parental lines were evaluated in a rain-out shelter (15 m wide x 75 m long) under wet and dry conditions in 2019 at the WWAI (-35.04540035261563, 147.34480619057837). The experiment was conducted in two adjacent blocks, wet and dry, where both blocks were managed with the same protocol except for watering. Within each block, plots were arranged in rectangular arrays with 24 rows and 9 ranges. The treatments comprised the factorial combinations of irrigation regimes: wet (control) and dry (stress); and lines. The treatments were allocated to plots in such a way that each irrigation regime was allocated to all plots in a single block (for operational convenience) and lines were allocated to plots within blocks. There were three replicate plots of all lines per block.

A total of 25 seeds were sown in single row-plots (1 m length) with custom-built stand, ensuring proper spacing between plants. Initially, a pre-seeding irrigation of 16 mm of water was applied across both blocks for optimal and uniform germination. Water stress treatment was applied at the stem elongation stage (BBCH scale 50) to the dry block, while stress was not applied to the wet block. Soil moisture was tracked on-line throughout the experiments using 1.5 m moisture probes inserted in ground (ICT) and ensured that plants were subjected to water stress. Three moisture probes were installed in the experiment following the manufacturer’s instructions: two in the dry block and one in the wet block; these probes were evenly spaced for reliable estimates of soil moisture across the blocks.

**Statistical methods**

(i) **QTL analysis for the Experiments 1-3**

The investigation of various traits on multiple trials required a multi-environment trial (MET) analysis for each trait. The MET data for each trait comprised the two field experiments (Experiments 1-2) and the pot experiment (Experiment 3) where each trial is considered as an “environment”. A baseline linear mixed model (LMM) is formulated first and then it is extended to a factor analysis form for the genotype by environment (GE) effects between environments. The GE effects are then partitioned into additive and non-additive GE effects by incorporating the marker data into the model. A working model for QTL analysis is then developed and genome scan is performed on this model to identify genomic regions associated with each trait. The potential set of markers identified from the genome scan is then thinned in order to establish a final multi-QTL model.

***Formulation of the baseline model***

The baseline models for each trait are presented here using the symbolic model notation of Wilkinson and Rogers (1973) and *ASReml-R* (Butler et al., 2018) syntax.

Δ^13^C

```
fixed ~  1 + Environment + Environment:Gdrop
random ~ diag(Environment):Gkeep +
at(Environment):Run +
               at(Environment):Carousal +
               at(Environment):Run:Carousal +
               at(Environment):Carousal:Tube +
               at(Environment):Block +
               at(Environment):Block:Plot
```

Days to flower

```
fixed ~  1 + Environment + Environment:Gdrop
random ~ diag(Environment):Gkeep +
at(Environment):Block +
at(Environment):Row +
               at(Environment):Range
```

Plant height

```
fixed ~  1 + Environment + Environment:Gdrop
random ~ diag(Environment):Gkeep +
at(Environment):Block +
               at(Environment):Block:Plot +
               at(Environment):Row +
               at(Environment):Range
```

Seed yield

```
fixed ~  1 + Environment + Environment:Gdrop
random ~ diag(Environment):Gkeep +
at(Environment):Block +
               at(Environment):Row +
               at(Environment):Range
```

NDVI

Fixed ~ 1 + Gdrop

random ~ Gkeep + Block + Row + Range

There are many characteristics of the above symbolic model notation that require explanation and defining. Firstly, formulation of the baseline model commenced with fitting a model that assumed independence of the GE effects between environments. This model termed the *diagonal (DIAG)* variance model for the GE effects (diag(Environment):Gkeep, where Gkeep is the lines that were genotyped) is analogous to analysing each environment separately. This baseline model is used to assess whether additional terms are required to account for non-treatment sources of variation as well as investigating the presence of outlier observations. Two additional terms were fitted as fixed effects, representing the main effects of the environments (Environment) and the interaction of the environments with those lines which were not genotyped but had phenotypic data (Environment:Gdrop, where Gdrop is the lines that were not genotyped) (see Tolhurst et al. (2019) for details). All models also included a term representing the overall mean (1) as a fixed effect.

The non-genetic component of the baseline model required the inclusion of random terms, which represent the plot structure of the randomized complete block designs of the experiments (Block), as well as accounting for other significant sources of non-genetic variation which occurred in either field/pot experiments (random Row or Range terms) or in the lab phase (see Table S3) for the trait Δ^13^C (Run, Carousal, Run:Carousal and Carousal:Tube) (see Smith et al. (2006) and Gilmour et al. (1997) for example). The working model also included terms (Block:Plot) which partition residual variance for those traits where the experimental unit is not the observational unit (see Bailey (2008)). These traits were plant height and Δ^13^C. The term at(Environment) denotes that variance models for these random model terms allowed for variance heterogeneity between environments. The variance models for the residuals were either independent and identically distributed (iid) effects (for Δ^13^C and plant height), or a two dimensional separable first order autoregressive variance model as explained in Cullis and Gleeson (1991) (for days to flower, seed yield and NDVI) with different variance parameters for each environment.

***Factor analytic variance models for the GE effects***

The baseline LMM for all traits except NDVI were then extended to include factor analytic (FA) variance models for the GE effects. A MET analysis was not necessary for NDVI as this trait was measured only in one environment (Experiment 2). The FA-LMM estimates the genetic variances and covariances between environments using a small number of unknown factors. These models were fitted using the reduced rank form of the FA model introduced by Thompson et al. (2003). Using the symbolic model notation of Wilkinson and Rogers (1973) and *ASReml-R* syntax, we fit

```
fixed ~ 1 + Environment + Environment:Gdrop
random ~ rr(Environment, k): Gkeep +
   diag(Environment): Gkeep
   …
```

where rr component models the common genotype by environment (CGE) effect, diag component models the specific genotype by environment (SGE) effect, k is the order of the model representing the number of unknown factors and “…” denotes other non-genetic terms. The FA modelling process commences with one factor (k = 1) and continues until either the limit of the data is reached or the overall percentage variance accounted for reaches 80%. For example, the limit for a MET dataset with $p = 3$ environments is k = 1 factor because an increase in the number of factors will result in more parameters being estimated than are possible in the fully unstructured model $(p(p + 1)/2= 6)$. For the FA1 model, there are $pk + p- k(k- 1)/2 = 6$ parameters to estimate.

In our case, an FA model of order 1 (FA1) was fitted for all traits except for seed yield, for which the variance of the SGE effects of one of the environments was constraint to be zero following Cullis & Smith (2016) as there were only two environments for this trait (Experiments 1-2). We refer to these models as baseline FA-LMM. On average, the percentage of genetic variation (%VAF) accounted by the baseline FA-LMM were above 86% for all traits (Δ^13^C: 88.6%, seed yield: 86.6%, days to flower: 92.3% and plant height: 89.9%). Estimated genetic correlations between environments were all greater than 0.83 for all traits.

***Partitioning GE effects into additive and non-additive GE effects***

The baseline FA-LMM was then extended to include marker information in order to determine the genomic regions that influence the traits associated with WUE. GE effects were partitioned into additive and non-additive GE effects (Oakey et al., 2007) and each were modelled by separate FA variance models. We utilised a genetic linkage map of 5101 DH population based on 1793 ‘bin’ DArTseq markers representing all 19 chromosomes of *B. napus*. The linkage map was constructed using the package *ASMap* (Taylor and Butler, 2014) in *R* statistical computing environment (R Core Team, 2019) utilising the Minimum spanning tree algorithm (Wu et al. 2008). The missing values in these markers were imputed using the *k*-nearest neighbour method (Troyanskaya et al., 2001). The resulting marker matrix, $M$, is used to construct a genomic relationship matrix, $K$, (GRM) (VanRaden, 2008) where $K=MM^{T}$ using the package *pedicure* (Butler, 2019) in *R* statistical computing environment (R Core Team, 2019). Using the symbolic model notation of Wilkinson and Rogers (1973) and *ASReml-R* syntax, we fit

```
fixed ~ 1 + Environment + Environment:Gdrop
random ~ rr(Environment, 1):vm(Gkeep, K)+
   diag(Environment):vm(Gkeep, K) +
   rr(Environment, 1):ide(Gkeep) +
   diag(Environment):ide(Gkeep) +
   …
```

where rr(Environment, 1):vm(Gkeep, K) models the additive CGE effects, diag(Environment):vm(Gkeep, K) models the additive SGE effects, rr(Environment, 1):ide(Gkeep) and diag(Environment):ide(Gkeep) models the same but for non-additive effects, K is the GRM and associated with the additive effects and “…” denotes other non-genetic terms. This model is equivalent to the standard genomic best linear unbiased prediction (GBLUP) model used in models for genomic selection (Tolhurst et al., 2019). We refer to this model as baseline FA-LMM with markers.

Both the between environment variance matrices of the GE additive genetic effects and non-additive genetic effects were modeled by FA structures of order 1 (FA1:FA1) for all traits except for seed yield, for which the variance of the additive and non-additive SGE effects of one of the environments was constraint to be zero as before. Table 1 presents a summary for baseline LMM, baseline FA-LMM and the baseline FA-LMM with markers in terms of the variance model for the GE effects, number of estimated parameters (total and genetic), residual log-likelihood and Akaike Information Criterion (AIC). The AIC value of the baseline FA-LMM with markers (or baseline LMM for NDVI) is the lowest for each trait, indicating that this model is the better fit.

Summary of the baseline models fitted for each trait with the variance model for genotype by environment (GE) effects, number of parameters estimated in each model in total (Total) and for the genetic variance (Genetic), log-likelihood (LogLik) and Akaike Information Criterion (AIC). DIAG: Diagonal; FA1: Factor analytic structure of order 1; FA1:FA1: Factor analytic structures of order 1 for both the between environment variance matrices of the GE additive genetic effects and non-additive genetic effects.

| **Model** | **Variance model for GE effects** | **Parameters** | | **LogLik** | **AIC** |
| --- | --- | --- | --- | --- | --- |
|  |  | **Total** | **Genetic** |  |  |
| *Δ^13^C* | | | | | |
| Baseline LMM | DIAG | 24 | 3 | -28.13016 | 104.2603 |
| Baseline FA-LMM | FA1 | 30 | 6 | 92.26117 | -124.5223 |
| Baseline FA-LMM with markers | FA1:FA1 | 39 | 12 | 112.21808 | -146.4362 |
| *Days to flower* | | | | | |
| Baseline LMM | DIAG | 17 | 3 | -3364.616 | 6763.232 |
| Baseline FA-LMM | FA1 | 23 | 6 | -2985.602 | 6017.204 |
| Baseline FA-LMM with markers | FA1:FA1 | 32 | 12 | -2888.462 | 5840.924 |
| *Plant height* | | | | | |
| Baseline LMM | DIAG | 18 | 3 | -23051.01 | 46138.02 |
| Baseline FA-LMM | FA1 | 24 | 6 | -22853.79 | 45755.58 |
| Baseline FA-LMM with markers | FA1:FA1 | 33 | 12 | -22837.59 | 45741.19 |
| *Seed yield* | | | | | |
| Baseline LMM | DIAG | 13 | 2 | 1110.801 | -2195.601 |
| Baseline FA-LMM | FA1 | 16 | 3 | 1244.776 | -2457.552 |
| Baseline FA-LMM with markers | FA1:FA1 | 21 | 6 | 1265.601 | -2489.202 |
| *NDVI* | | | | | |
| Baseline LMM | - | 7 | 1 | 938.9046 | -1863.809 |
| Baseline LMM with markers | - | 8 | 2 | 943.0174 | -1870.035 |

***Working model for QTL analysis***

The approach used herein for the determination of genomic regions that influence the expression of the traits associated with water use efficiency in multiple environments is an extension of the whole genome, single-step, multi-environment QTL analysis approach developed by Verbyla et al. (2012).

Our approach uses an alternative working model where we partition $M$ as $M=\left[ M_{1} M_{2}\ldots M_{c} \right]$ and let $K= \sum_{i=1}^{c} M_{i}M_{i}^{T}$. Further, let $M_{i}=\left( m_{i;jk} \right)$ be the marker matrix for the chromosome $i$ of size $n_{g}$× $n_{m_{i}}$ where $n_{g}$is the number of genotypes with marker data, $n_{m_{i}}$is the number of markers in chromosome $i$ and $j$ and $k$ are the subscripts for markers within chromosomes and genotypes respectively. If $n_{m}= \sum_{i=1}^{c} n_{m_{i}}$ then $M$ is $n_{g}$× $n_{m}$ and $K$ is $n_{g}$× $n_{g}$.

Let baseline FA-LMM with markers denoted by FA_B_. For the QTL analysis we fit the same FA_B_ model where we replace the GRM, $K$, by the formulation presented above. Further, let $\hat{\kappa}^{T}=({\hat{\kappa}^{T}}_{vm}{, \hat{\kappa}^{T}}_{ide}, {\hat{\kappa}^{T}}_{o})$ be the vector of variance parameters partitioned conformably with the associated model terms and evaluated at the residual maximum likelihood (REML) estimates of $\hat{\kappa}$ from FA_B_.

The QTL analysis approach that we use has multiple steps which are described in detail in the following.

**Step 1: Genome scan**

To avoid proximal contamination (Listgarten et al. 2012) and to reduce computation time (Lippert et al. 2011) we scan the genome by fitting each marker $m_{i;j}$, $i=1,\ldots, c$ and $j=1, \ldots, n_{m_{i}}$ as a covariate by adding the main effect and the interaction of the marker within the environment. In this fit we reduce computation by holding $\kappa_{vm}$ at the REML estimates from FA_B_ and only re-estimate $\kappa_{ide}$ and $\kappa_{o}$. Furthermore, we replace the GRM, $K$, by $K_{-i}$ where $K_{-i}$ is given by

$$K_{-i}= \sum_{\begin{aligned} l=1, \\ l\neq i \end{aligned}}^{c} M_{l}M_{l}^{T}$$

This avoids the possibility of proximal contamination and is a conservative approach.

For each fit we compute the preceding terms’ Wald statistics from the fit of

fixed ~ Environment + Environment:Gdrop + $m_{i;j}$ + $m_{i;j}$:Environment

where $m_{i;j}$ represents the covariate $m_{i;j}$ expanded by $m_{i;j}=Z_{g}m_{i;j}$ where $Z_{g}$ is the design matrix for Gkeep. Two probabilities are obtained for each fit and we denote these by $p_{m;ij}$ and $p_{m;ij.e}$

**Step 2: LD block thinning for Marker by Environment (**${\boldsymbol{M}\boldsymbol{\times}\boldsymbol{E}}_{\boldsymbol{Q}_{\boldsymbol{ME}}}$**) interactions**

In this step, we determined LD blocks by calculating the squared LD ($r^{2}$) for each pair of markers and any markers with an estimated LD of greater than 0.7 were deemed to be within the same LD block. We thin ${\{p}_{m;ij.e}\}$ by tagging each $p_{m;ij.e}$ by its LD block and choosing the representative marker to be the marker with the lowest *p*-value from the set of markers in each LD block. If we denote the set of thinned *p-*values by ${p_{me}}^{T}$, we then apply a false discovery rate (FDR) approach (Benjamini and Hochberg, 1995) to this vector to determine the working set of Marker by Environment (${M\times E}_{Q_{ME}}$) QTLs.

**Step 3: LD block thinning for Marker (**$\boldsymbol{M}_{\boldsymbol{Q}_{\boldsymbol{M}}}$**) main effects**

In this step, we apply the same approach from Step 2 to thin ${\{p}_{m;ij}\}$, except before thinning we discard any marker $m_{i;j}$ that is present in the thinned set from Step 2, i.e. in the working set of ${M\times E}_{Q_{ME}}$ QTLs. We also exclude all markers from the set of LD blocks associated with the working set of ${M\times E}_{Q_{ME}}$ QTLs. This provides the working set denoted by $M_{Q_{M}}$.

**Step 4: Final multi QTL model**

We include all markers from Step 2 ${M\times E}_{Q_{ME}}$ QTL set as “$m_{i;j}+ m_{i;j}$:Environment” and include all markers from Step 3 $M_{Q_{M}}$ QTL set as “$m_{i;j}$” as fixed terms in the model. In this model we replace the GRM, $K$, by $K_{e}$, where $K_{e}$ is the GRM using markers from LD blocks which do not contain markers from working QTL sets $M_{Q_{M}}$and ${M\times E}_{Q_{ME}}$. We then use standard backward elimination procedure to determine the final multi-QTL model. During this procedure we remove the ${M\times E}_{Q_{ME}}$ interactions first and foremost and stop when there are no more non-significant interactions. We then focus on the $M_{Q_{M}}$main effects but leaving the main effects for the significant interactions in the model, i.e. only exclude main effects for those non-significant interactions and then exclude main effects only if there is not a partner interaction. Each time when a marker is dropped during the backward elimination, that marker and its LD block is added back into the GRM. All remaining $M_{Q_{M}}$ and ${M\times E}_{Q_{ME}}$ markers in the fixed component of the final multi-QTL model are reported as putative QTLs together with their positions, LOD scores, i.e., –log_10_(*p*-value) and the percentage of genetic variance accounted ($R^{2}$).

(ii) **LMM analysis for the Experiment 4**

The models for each trait for the LMM analysis of Experiment 4 are presented here using the symbolic model notation of Wilkinson and Rogers (1973).

Δ^13^C

fixed ~ 1

random ~ Genotype + Carousel + Rep + Plot

Days to flower

fixed ~ 1

random ~ Genotype + Block:Rep + Block:Range + Block:Row

Plant height

```
fixed ~ 1 + Irrig[Block]
random ~ Genotype + Genotype:Irrig[Block] + Block[Irrig]:Rep  + Plot + Block[Irrig]:Range + Block[Irrig]:Row
```

Seed yield

```
fixed ~ 1 + Irrig[Block] + nplants
random ~ Genotype + Genotype:Irrig[Block] + Block[Irrig]:Rep  + Block[Irrig]:Range + Block[Irrig]:Row
```

Gas exchange and physiological traits

fixed ~ 1

random ~ Genotype + Rep + Range + Row

There are many characteristics of the above symbolic model notation that require explanation and defining. Firstly, the construction of the appropriate genetic component of the model for each trait represented the treatment structure of the trial. As there was a factorial treatment structure (factorial combination of genotypes and irrigation regimes), it is necessary to incorporate this structure in the analysis. Based on the randomisation procedure of the factorial treatment structure described in Method S1, there is aliasing of irrigation regime (Irrig) and block (Block) effects and hereafter denoted as Irrig[Block] or Block[Irrig]. Hence, there was no valid inferential framework to test the main effect of irrigation regimes (Irrig). This is explained as “pseudo” or “false" replication in Bailey (2008). There was a valid inferential framework to test the genotypes by irrigation regimes interaction (Genotype:Irrig[Block]), but caution should be exercised when interpreting the results as these effects may be due to the different blocks (Block[Irrig]) (see Gururaj et al., (2020) for details). Thus, in terms of the treatment effects, random genotype main effects (Genotype) and random genotype by irrigation block interactions were fitted (Genotype:Irrig[Block]) and the Irrig[Block] effects were fitted as fixed. However, this treatment structure is only appropriate for the traits plant height and seed yield as days to flower is measured before the irrigation regime treatment being imposed, whereas Δ^13^C, gas exchange and physiological measurements were taken only for the wet block. Hence, for the remaining traits, only the genotype (Genotype) main effects were fitted as random. The seed yield is measured for each plot and the number of plants (nplants) available in each plot were measured. Number of plants is fitted as a covariate accounting for the variation in seed yield (*p*-value = 0.000057507 from Wald test). All models also included a term representing the overall mean (1) as a fixed effect.

The non-genetic component of the model for each trait required the inclusion of random terms representing the plot structure of the experiment (Block and Block:Rep). For plant height and seed yield random block (Block) effects were not fitted as the aliasing irrigation regime effects (Irrig[Block]) were fitted as fixed. As Δ^13^C, gas exchange and physiological measurements were taken only for the wet block these traits required inclusion of only the random Rep effects. Random row (Block:Row or Row) and range (Block:Range or Range) effects within blocks were included as necessary. For plant height and Δ^13^C, the model also included random Plot effects which partition residual variance as the experimental unit of these traits is not the observational unit. For Δ^13^C, random Carousel effects were fitted as it is a source of non-genetic variation which occurred in the lab phase. Variations related to the runs were not necessary as all samples were evaluated in a single run (see Table S3). The variance models for the residuals were independent and identically distributed (iid) effects as the experiment was conducted in single row plots.

***Light microscopy***

A leaf disc (9.08 cm^2^ size) was taken from each of two replicate canola lines from Experiment 4 (wet treatment) for anatomical analysis. Discs were collected from leaf blades, at a position between the 2nd and 3rd veins, from plants at the flowering stage and fixed in 70% ethanol. After clearing in 5% household bleach (~0.2% sodium hypochlorite) for five days, they were stored in 70% ethanol before sectioning and staining using a method modified from Rae *et al.* (2020). For imaging, hand sections were made from leaf discs, oxidised in 1% (w/v) periodic acid for ~15 sec, and then rinsed in water ~15 sec. Sections were incubated in 2 µg/ml Rhodamine-123 in water for 30 min and rinsed for 5-10 min in water before mounting in 50% glycerol. Samples were imaged using 488 nm excitation and 500-560 nm emission on a Leica SP8 confocal microscope.

RNA sequencing and differential gene expression analysis

Parental lines, BC1329 and BC9102 of DH population were grown in three replicates under both wet (100% field capacity) and dry (50% field capacity) treatments in a glasshouse, located at Wagga Wagga Agricultural Institute . The clean sequence reads (100 bp single-end reads) for 12 samples that had per base sequence quality with >96% bases above Q30 were aligned against the *B. napus* reference Darmor-*bzh* (Version 4.1), using STAR aligner (v2.5.3a) (https://github.com/alexdobin/STAR/blob/master/doc/STARmanual.pdf). The raw counts of reads mapping to each known gene was used to perform differential expression analysis using edgeR (version 3.30.3) (https://bioconductor.org/packages/release/bioc/html/edgeR.html) using R version 4.0.3. Counts are summarised at gene level using the featureCounts v1.5.3 utility of the subread package (<http://subread.sourceforge.net/>). The default TMM normalisation method of edeR was used to normalise the counts between samples. A generalised linear model approach was then used to quantify the differential expression between the groups. The differentially expressed genes (DEGs) were obtained using a false discovery rate (FDR < 0.05). Heatmaps showing the expression pattern of genes in A09 and C09 QTL regions were produced using the ComplexHeatmap R package (Gu et al., 2016).

**References support Statistical analysis**

**Bailey, R. (2008).** *Design of Comparative Experiments (Cambridge Series in Statistical and Probabilistic Mathematics).* Cambridge: Cambridge University Press. doi: 10.1017/CBO9780511611483

**Benjamini, Y. and Hochberg, Y. (1995).** Controlling the false discovery rate: a practical and powerful approach to multiple testing. *Journal of the Royal Statistical Society: Series B* 57, 289–300. doi: 10.1111/j.2517-6161.1995.tb02031.x

**Butler, D. G., Cullis, B. R., Gilmour, A. R., Gogel, B. J. and Thompson, R. (2018).** ASReml-R Reference Manual Version 4. Technical report, VSN International Ltd, Hemel Hempstead, HP1 1ES, UK.

**Cullis, B. R. and Gleeson, A. (1991).** Spatial Analysis of Field Experiments-An Extension to Two Dimensions. *Biometrics*, 47(4), 1449-1460. doi:10.2307/2532398

**Cullis, B. R. and Smith, A. B. (2016).** The analysis of QTL and QTL by treatment experiments using spatial models for marker effects. Technical report.

**Gilmour, A. R., Cullis, B. R. and Verbyla, A. P. (1997).** Accounting for natural and extraneous variation in the analysis of field experiments. *Journal of Agricultural, Biological, and Environmental Statistics* 2, 269–293. doi: 10.2307/1400446

**Kadkol, G., Smith, A., Cullis, B. and Chenu, K. (2020).** Variation in Australian durum wheat germplasm for productivity traits under irrigated and rainfed conditions: Genotype performance for agronomic traits and benchmarking. *The Journal of Agricultural Science* 158(6), 479-495. doi:10.1017/S0021859620000817

**Lippert, C., Listgarten, J., Liu, Y., Kadie, C. M., Davidson, R. I. and Heckerman D (2011**). FaST linear mixed models for genome-wide association Studies. *Nature methods*, 8, 833–835. https://doi.org/10.1038/nmeth.1681

**Listgarten, J., Lippert, C., Kadie, C. M., Davidson, R. I., Eskin, E. and Heckerman, D. (2012).** Improved linear mixed models for genome-wide association studies. *Nature methods*, 9(6), 525–526.

**Oakey, H., Verbyla, A., Cullis, B. R., Wei, X. and Pitchford, W. (2007).** Joint modelling of additive and non-additive (genetic line) efects in multi-environment trials. *Theoretical and Applied Genetics* 114, 1319–1332.

**Raman, H., McVittie, B., Pirathiban, R., Raman, R., Zhang, Y., Barbulescu, DM., Qiu, Y., Liu, S. and Cullis B. (2020).** Genome-Wide Association Mapping Identifies Novel Loci for Quantitative Resistance to Blackleg Disease in Canola. *Frontiers in Plant Science* 2020; 11:1184. pmid:32849733

**Smith, A., Lim, P. and Cullis, B. (2006).** The design and analysis of multi-phase plant breeding experiments. *The Journal of Agricultural Science* 144 5, 393–409. doi: 10.1017/ S0021859606006319

**Smith, A., Ganesalingam, A., Kuchel, H. and Cullis, B. (2015).** Factor analytic mixed models for the provision of grower information from national crop variety testing programmes. *Theoretical and Applied Genetics* 128:55–72

**Taylor, J. D. and Butler, D. (2014). ASMap:** An (A)ccurate and (S)peedy linkage map construction package for inbred populations that uses the extremely efficient MSTmap algorithm.

**Thompson, R., Cullis, B., Smith, A. and Gilmour, A. (2003).** A Sparse Implementation of the Average Information Algorithm for Factor Analytic and Reduced Rank Variance Models. *Australian and New Zealand Journal of Statistics* 45, 445-459.

**Tolhurst, D. J., Mathews, K. L., Smith, A. B. and Cullis, B. R. (2019).** Genomic selection in multi-environment plant breeding trials using a factor analytic linear mixed model. *Journal of Animal Breeding and Genetics* 136, 279–300. doi: 10.1111/ jbg.12404

**Troyanskaya, O., Cantor, M., Sherlock, G., Brown, P., Hastie, T., Tibshirani, R., et al. (2001).** Missing value estimation methods for DNA microarrays. *Bioinformatics* 17, 520–525. doi: 10.1093/bioinformatics/17.6.520

**VanRaden, P. M. (2008).** Efficient Methods to Compute Genomic Predictions. *Journal of Dairy Science,* 91, 4414–4423. https ://doi.org/10.3168/jds.2007-0980

**Verbyla, A.P. and Cullis, B.R.** Multivariate whole genome average interval mapping: QTL analysis for multiple traits and/or environments. *Theoretical and Applied Genetics* 125, 933–953 (2012). https://doi.org/10.1007/s00122-012-1884-9

**Wilkinson, G. N. and Rogers, C. E. (1973).** Symbolic description of factorial models for analysis of variance. *Journal of the Royal Statistical Society. Series C (Applied Statistics)* 22, 392–399.

**Wu, Y., Bhat, P. R., Close, T. J. and Lonardi, S. (2008).** Efficient and accurate construction of genetic linkage maps from the minimum spanning tree of a graph. *PLoS Genetics* 4.
