## Supplementary material for "Multi-environment QTL analysis delineates a major locus associated with homoeologous exchanges for water-use efficiency and seed yield in allopolyploid *Brassica napus*": Table S5

**Table S5:** Summary of the partitioning of genetic variance into Additive and Non-additive and REML estimate of the Total (additive plus non-additive) genetic variance before (Baseline FA-LMM with markers: M1) and after identifying putative QTL (Final multi QTL model: M2) for each of the trait and trial.

| **Trait** | **Experiment** | **Phenotypic environment** | **Additive (M1, %)** | **Additive (M2, %)** | **Non-additive (M1, %)** | **Non-additive (M1, %)** | **Total (M1)** | **Total (M2)** | **VAF_m_ (%)** |
| --- | --- | --- | --- | --- | --- | --- | --- | --- | --- |
| Δ^13^C | 1 | Field (2017) | 47.99 | 23.46 | 52.01 | 78.36 | 0.49 | 0.32 | - |
|  | 2 | Field (2018) | 50.11 | 23.98 | 49.89 | 77.88 | 0.23 | 0.15 | - |
|  | 3 | Pot (2017) | 55.65 | 19.96 | 44.35 | 81.59 | 0.23 | 0.12 | - |
|  |  | Mean | 51.25 | 22.47 | 48.75 | 79.28 | 0.32 | 0.20 | 37.91 |
| DTF | 1 | Field (2017) | 79.41 | 31.77 | 20.59 | 72.66 | 45.14 | 17.16 | - |
|  | 2 | Field (2018) | 82.44 | 34.07 | 17.56 | 70.68 | 55.66 | 19.07 | - |
|  | 3 | Pot (2017) | 75.55 | 13.19 | 24.45 | 88.64 | 46.02 | 17.08 | - |
|  |  | Mean | 79.13 | 26.35 | 20.87 | 77.33 | 48.94 | 17.77 | 63.68 |
| NDVI | 2 | Field (2018) | 21.53 | 13.92 | 78.47 | 86.08 | 0.01 | 0.01 | 11.76 |
| PH | 1 | Field (2017) | 19.04 | 1.92 | 80.96 | 98.22 | 135.09 | 108.43 | - |
|  | 2 | Field (2018) | 34.42 | 11.00 | 65.58 | 89.79 | 131.97 | 88.13 | - |
|  | 3 | Pot (2017) | 45.87 | 22.57 | 54.13 | 79.04 | 69.30 | 43.63 | - |
|  |  | Mean | 33.11 | 11.83 | 66.89 | 89.02 | 112.12 | 80.06 | 28.59 |
| SY | 1 | Field (2017) | 40.96 | 2.91 | 59.04 | 98.69 | 0.14 | 0.08 | - |
|  | 2 | Field (2018) | 60.18 | 17.99 | 39.82 | 91.92 | 0.08 | 0.04 | - |
|  |  | Mean | 50.57 | 10.45 | 49.43 | 95.31 | 0.11 | 0.06 | 45.42 |

VAF_m_ shows the percentage of genetic variance accounted by the identified putative QTLs. Trait variations were assessed using MET analyses across three environments in 2017 and 2018 under field and pot experiments (Experiments 1-3) for Δ^13^C, DTF and PH and two environments (Experiments 1-2) for SY; whereas it was based on a single site analysis from only one environment for NDVI (Experiment 2). Δ^13^C: Carbon isotope discrimination; DTF: Days to flower; NDVI: Normalised difference vegetative difference; PH: Plant height; SY: Seed yield.
