## Supplementary material for "Multi-environment QTL analysis delineates a major locus associated with homoeologous exchanges for water-use efficiency and seed yield in allopolyploid *Brassica napus*": Table S6

**Table S6:** Summary of heritability, mean, minimum and maximum values for each trait and trial in the doubled haploid population from BC1329/BC9102.

| **Trait** | **Experiment** | **Phenotypic environment** | **Mean** | **Minimum** | **Maximum** | **Heritability**  **(*h^2^*)** |
| --- | --- | --- | --- | --- | --- | --- |
| Δ^13^C | 1 | Field (2017) | 20.47 | 18.66 | 21.75 | 0.86 |
|  | 2 | Field (2018) | 18.87 | 17.59 | 19.86 | 0.59 |
|  | 3 | Pot (2017) | 21.21 | 20.10 | 22.08 | 0.59 |
|  |  | OP | 20.16 | 18.73 | 21.25 | - |
| DTF | 1 | Field (2017) | 116.1 | 101.3 | 131.5 | 0.98 |
|  | 2 | Field (2018) | 115.00 | 98.19 | 132.42 | 0.98 |
|  | 3 | Pot (2017) | 108.07 | 92.64 | 124.05 | 0.91 |
|  |  | OP | 113.07 | 97.37 | 129.33 | - |
| NDVI | 2 | Field (2018) | 0.4604 | 0.1734 | 0.6230 | 0.92 |
| PH | 1 | Field (2017) | 80.34 | 46.18 | 99.61 | 0.93 |
|  | 2 | Field (2018) | 89.26 | 53.34 | 109.52 | 0.91 |
|  | 3 | Pot (2017) | 77.84 | 53.73 | 91.45 | 0.56 |
|  |  | OP | 82.48 | 51.08 | 100.20 | - |
| SY | 1 | Field (2017) | 0.53 | 0.02 | 1.25 | 0.98 |
|  | 2 | Field (2018) | 0.41 | 0.001 | 0.99 | 0.98 |
|  |  | OP | 0.47 | 0.01 | 1.12 | - |

Trait variations were assessed using MET analyses across three environments in 2017 and 2018 under field and pot experiments (Experiments 1-3) for Δ^13^C, DTF and PH and two environments (Experiments 1-2) for SY; whereas it was based on a single site analysis from only one environment for NDVI (Experiment 2). CGE-EBLUPs and Overall performance (OP) estimates are summarised for Δ^13^C, DTF, PH and SY; whereas genotype EBLUPs are summarised for NDVI. Δ^13^C: Carbon isotope discrimination; DTF Days to flower; NDVI: Normalised difference vegetative difference; PH: Plant height; SY: Seed yield.
