## Supplementary material for "Multi-environment QTL analysis delineates a major locus associated with homoeologous exchanges for water-use efficiency and seed yield in allopolyploid *Brassica napus*": Table S7

**Table S7:** REML estimate of the Between environment genetic correlation matrices for Additive and Total (additive plus non-additive) effects of each trait from the baseline FA-LMM with markers.

| **Trait** | **Variance** | **Phenotypic environment** | **Correlation (ρ)** | | |
| --- | --- | --- | --- | --- | --- |
|  |  |  | Field (2017) | Field (2018) | Pot (2017) |
| Δ^13^C | Additive | Field (2017) | 1.00 | 0.96 | 0.99 |
|  |  | Field (2018) | 0.96 | 1.00 | 0.98 |
|  |  | Pot (2017) | 0.99 | 0.98 | 1.00 |
|  | Total | Field (2017) | 1.00 | 0.93 | 0.83 |
|  |  | Field (2018) | 0.93 | 1.00 | 0.87 |
|  |  | Pot (2017) | 0.83 | 0.87 | 1.00 |
| DTF | Additive | Field (2017) | 1.00 | 0.98 | 0.92 |
|  |  | Field (2018) | 0.98 | 1.00 | 0.94 |
|  |  | Pot (2017) | 0.92 | 0.94 | 1.00 |
|  | Total | Field (2017) | 1.00 | 0.94 | 0.91 |
|  |  | Field (2018) | 0.94 | 1.00 | 0.90 |
|  |  | Pot (2017) | 0.91 | 0.90 | 1.00 |
| PH | Additive | Field (2017) | 1.00 | 0.95 | 0.94 |
|  |  | Field (2018) | 0.95 | 1.00 | 0.89 |
|  |  | Pot (2017) | 0.94 | 0.89 | 1.00 |
|  | Total | Field (2017) | 1.00 | 0.90 | 0.87 |
|  |  | Field (2018) | 0.90 | 1.00 | 0.94 |
|  |  | Pot (2017) | 0.87 | 0.94 | 1.00 |
| SY | Additive | Field (2017) | 1.00 | 0.91 | - |
|  |  | Field (2018) | 0.91 | 1.00 | - |
|  | Total | Field (2017) | 1.00 | 0.87 | - |
|  |  | Field (2018) | 0.87 | 1.00 | - |

Trait variations were assessed using MET analyses across three environments in 2017 and 2018 under field and pot experiments (Experiments 1-3) for Δ^13^C, DTF and PH and two environments (Experiments 1-2) for SY. Δ^13^C: Carbon isotope discrimination; DTF: Days to flower; PH: Plant height; SY: Seed yield; -: Not applicable.
