## Supplementary material for "Multi-environment QTL analysis delineates a major locus associated with homoeologous exchanges for water-use efficiency and seed yield in allopolyploid *Brassica napus*": Table S9

**Table S9**. **Summary of heritability, mean, minimum and maximum values of the genotype EBLUPs for each trait evaluated in the rain-out shelter (Experiment 4) with wet and dry conditions for a doubled haploid population from BC1329/BC9102.**

| ***Trait** | **Units** | **Block** | **Heritability (*H*^2^)** | **Mean** | **Minimum** | **Maximum** |
| --- | --- | --- | --- | --- | --- | --- |
| Δ^13^C | ‰ | Wet | 0.39 | 22.91 | 22.17 | 23.49 |
| *A* | µmol m^-2^ sec^-1^ | Wet | 0.77 | 10.34 | 4.97 | 17.15 |
| *g_sw_* | mol H_2_O m^-2^ sec^-1^ | Wet | 0.58 | 0.23 | 0.11 | 0.38 |
| *i*WUE | µmol/ mol H_2_O | Wet | 0.31 | 45.68 | 30.89 | 60.41 |
| LWC | - | Wet | 0.42 | 10.14 | 8.36 | 12.52 |
| DTF | days | Wet | 0.97 | 107.96 | 84.20 | 132.41 |
| PH | cm | Dry | 0.93 | 157.2 | 122.6 | 195.6 |
|  |  | Wet |  | 157.3 | 120.0 | 193.0 |
| SY | g | Dry | 0.89 | 60.45 | 10.81 | 108.63 |
|  |  | Wet |  | 110.47 | 62.76 | 163.50 |

*****Δ^13^C: Carbon isotope discrimination; *A*: Photosynthesis rate; g_sw_: stomatal conductance; SLW: iWUE: intrinsic water use efficiency; LWC: Leaf water content; DTF: Days to flower; PH: Plant height; SY: Seed yield. Seed yield is predicted at the average value of 9.2 plants per plot.
