## Supplementary material for "Multi-environment QTL analysis delineates a major locus associated with homoeologous exchanges for water-use efficiency and seed yield in allopolyploid *Brassica napus*": Fig S1

Fig. S1. Weather conditions prevailed under field conditions (Wagga Wagga, NSW 2650, Australia), where experiments were conducted during canola growing seasons (May to November) in 2017, and 2018. a: Mean monthly maximum and minimum temperatures, b: Monthly cumulative total rainfall.

a

Seed Sowing	Seed harvesting

Mean Monthly Temperature (oC)

| Month | 2017 Max | 2018 Max | 2017 Min | 2018 Min |
| --- | --- | --- | --- | --- |
| Jan | 35.1 | 34.3 | 17.6 | 18.3 |
| Feb | 32.7 | 32.4 | 15.6 | 16.2 |
| Mar | 30.2 | 29.9 | 15.9 | 13.6 |
| Apr | 23.1 | 27.7 | 9.4 | 11.2 |
| May | 18.1 | 18.4 | 5.3 | 5.8 |
| Jun | 15.1 | 14.8 | -0.1 | 4.1 |
| Jul | 13.7 | 13.8 | 2.6 | 1.7 |
| Aug | 14.5 | 14.7 | 2.6 | 2.9 |
| Sep | 19.3 | 19.1 | 4.9 | 3.8 |
| Oct | 24.3 | 26 | 9.2 | 9.5 |
| Nov | 28 | 26.3 | 13.9 | 13 |
| Dec | 30.5 | 32.2 | 16.6 | 17.5 |

Seed Sowing	Seed harvesting

b

Cumulative Total Rainfall (mm)

| Month | 2017 | 2018 |
| --- | --- | --- |
| Jan | 13.8 | 50.8 |
| Feb | 31.8 | 75 |
| Mar | 76.6 | 82.6 |
| Apr | 109 | 86.6 |
| May | 129.2 | 122 |
| Jun | 131.4 | 162.8 |
| Jul | 184 | 177 |
| Aug | 233.7 | 198.8 |
| Sep | 242.1 | 231 |
| Oct | 307.5 | 252 |
| Nov | 343.7 | 347.6 |
| Dec | 445.3 | 404.2 |
