## Supplementary material for "Multi-environment QTL analysis delineates a major locus associated with homoeologous exchanges for water-use efficiency and seed yield in allopolyploid *Brassica napus*": Fig S2

Fig. S2. Layout of experiments conducted under field in 2017 (**A**, Experiment 1), 2018 (**B**, Experiment 2), and under rain-out shelter conditions in 2017 (**C**, Experiment 3) and 2019 (**D**, Experiment 4).

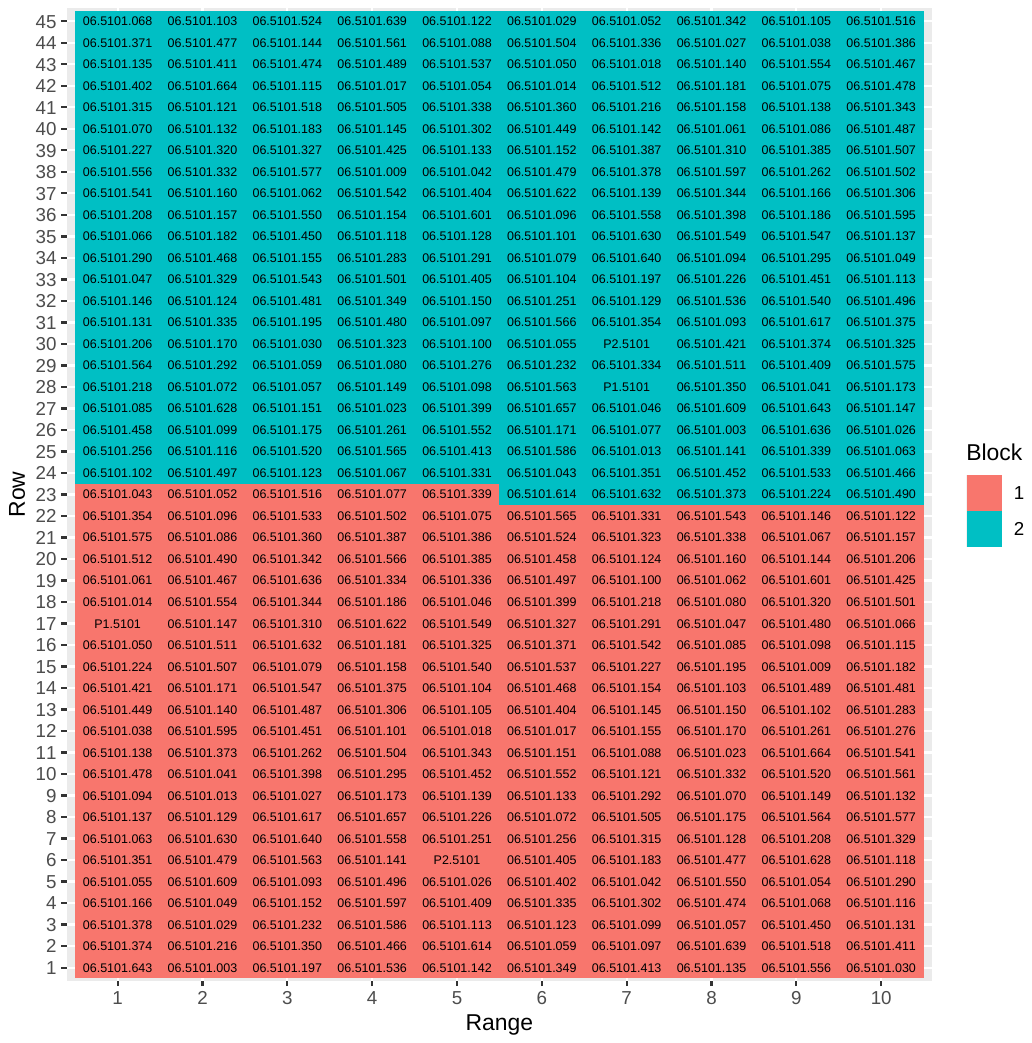
**A**

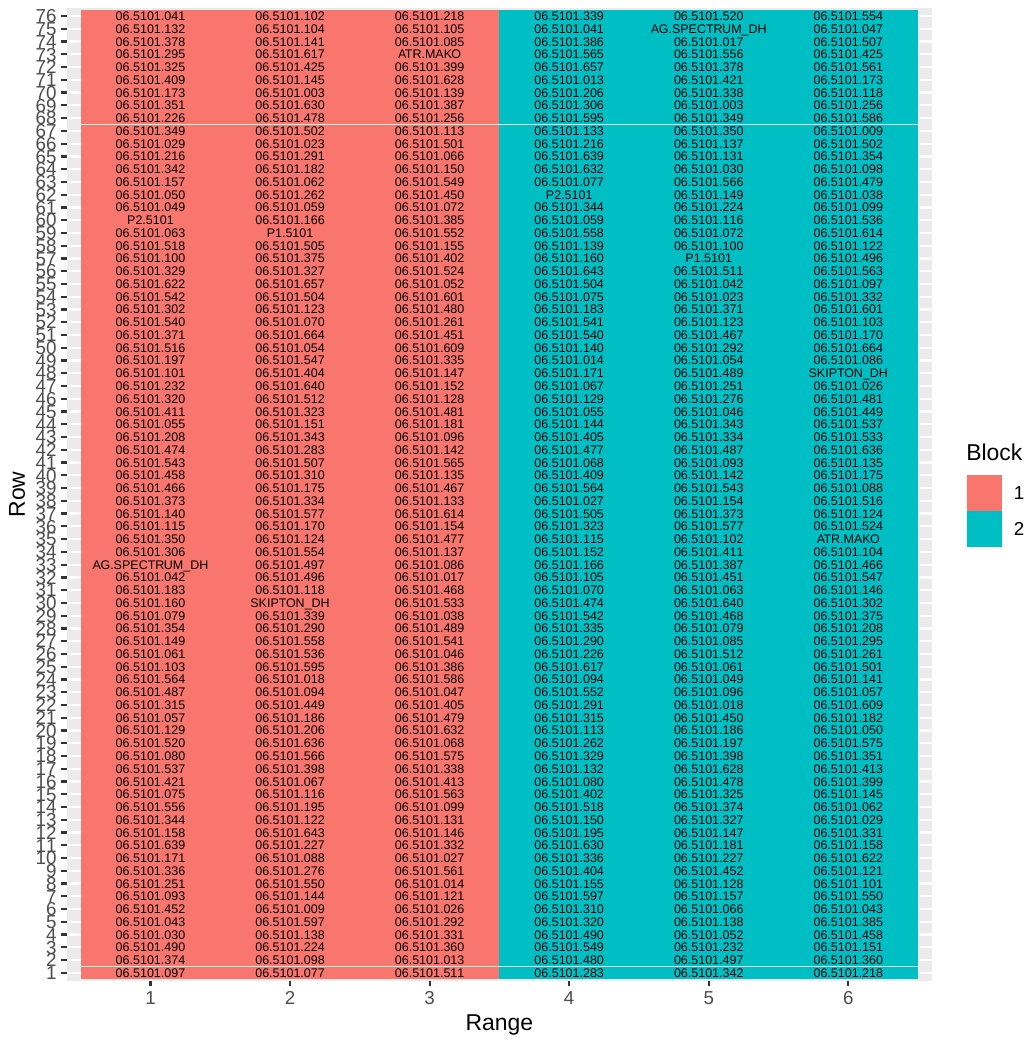
**B**

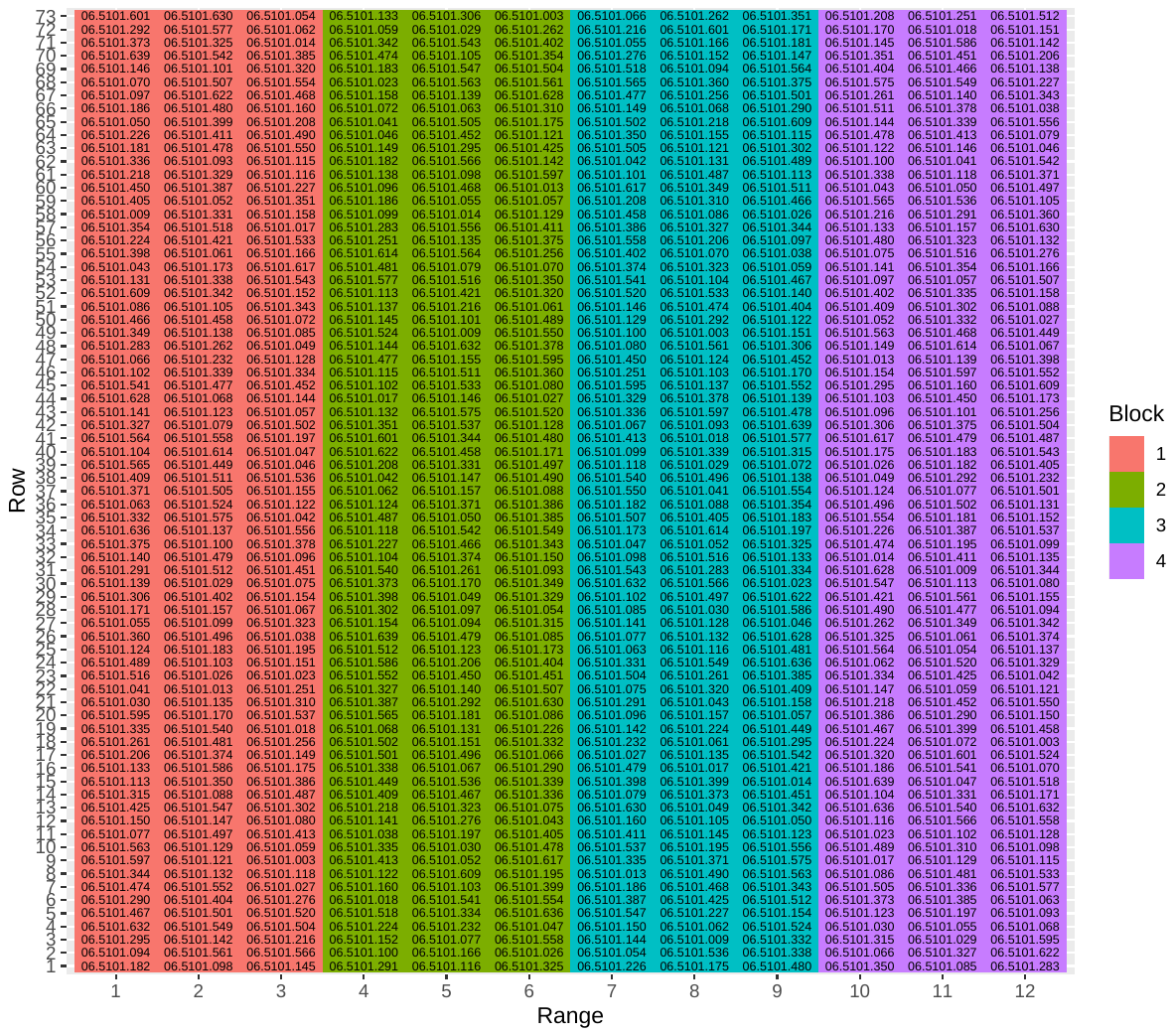
**C**

**D**

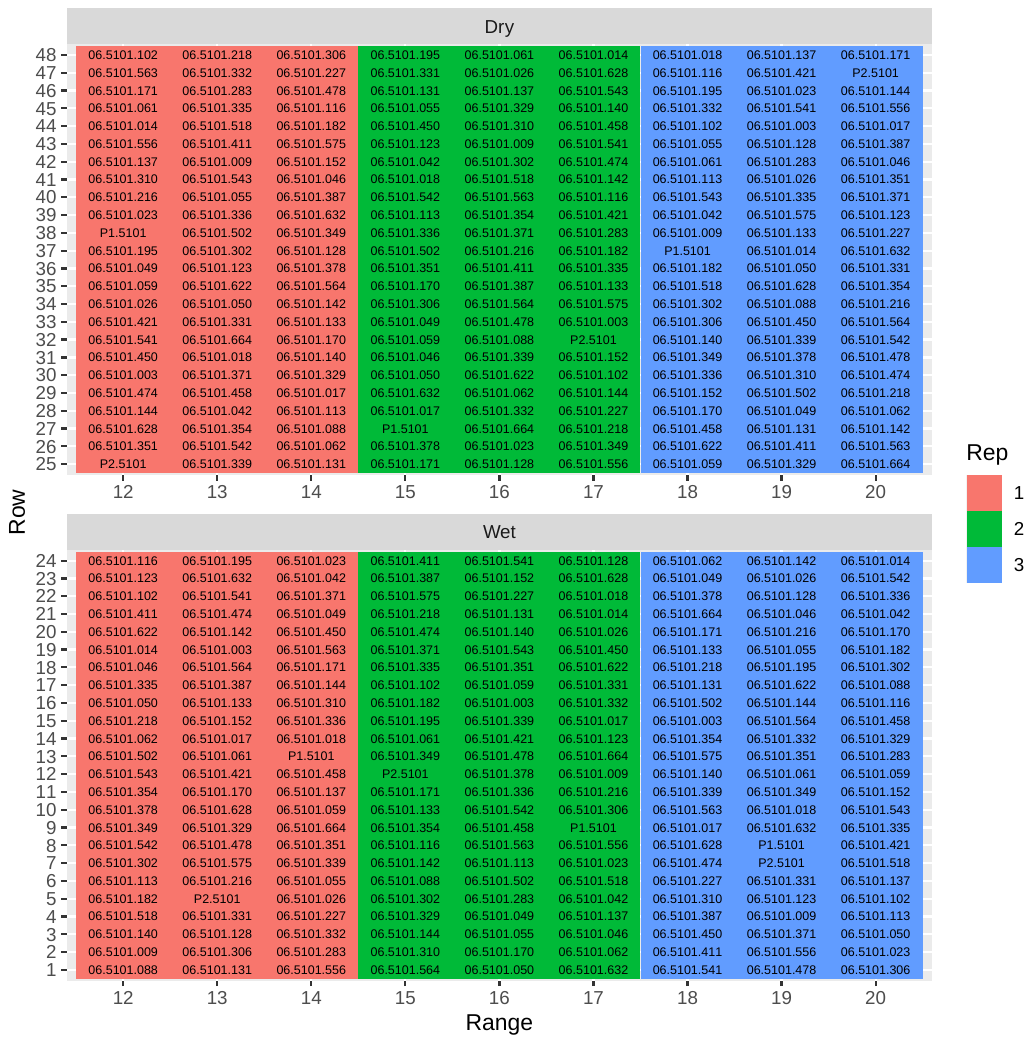
