## Supplementary material for "Multi-environment QTL analysis delineates a major locus associated with homoeologous exchanges for water-use efficiency and seed yield in allopolyploid *Brassica napus*": Fig S3

Fig. S3 Graphical representation showing localisation of multi-trait QTL associated with Δ^13^C (‰), flowering time (days to flower, DTF); plant height (PHT) and seed yield (SY) on chromosomes A09 (a) and C09 (b) in a DH population from the BC1329/BC9102. DArTseq markers and their genetic map positions are shown on *right*- and *left*-hand side, respectively. Solid lines (in blue and red colour) represent markers that showed significant associations with traits of interest. Map distances are given in cM and displayed using the MapChart.

a

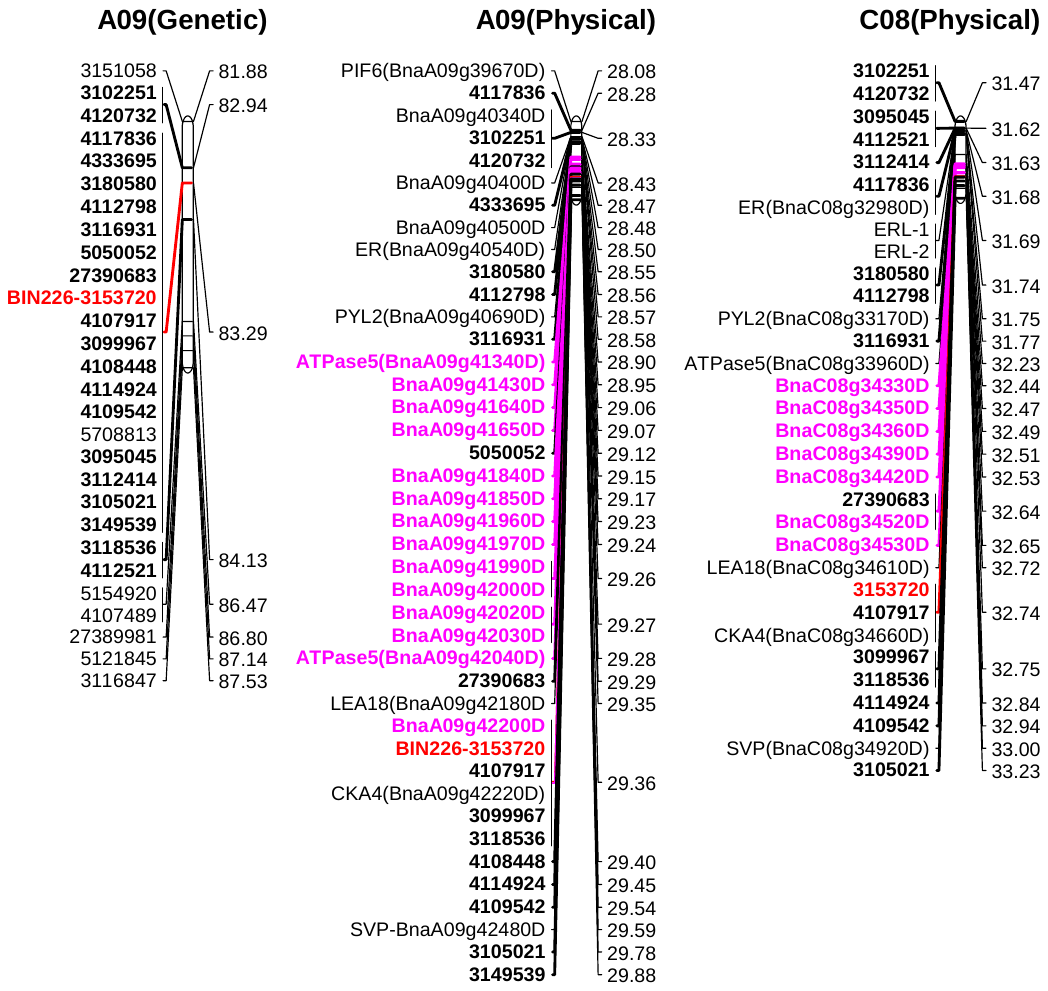

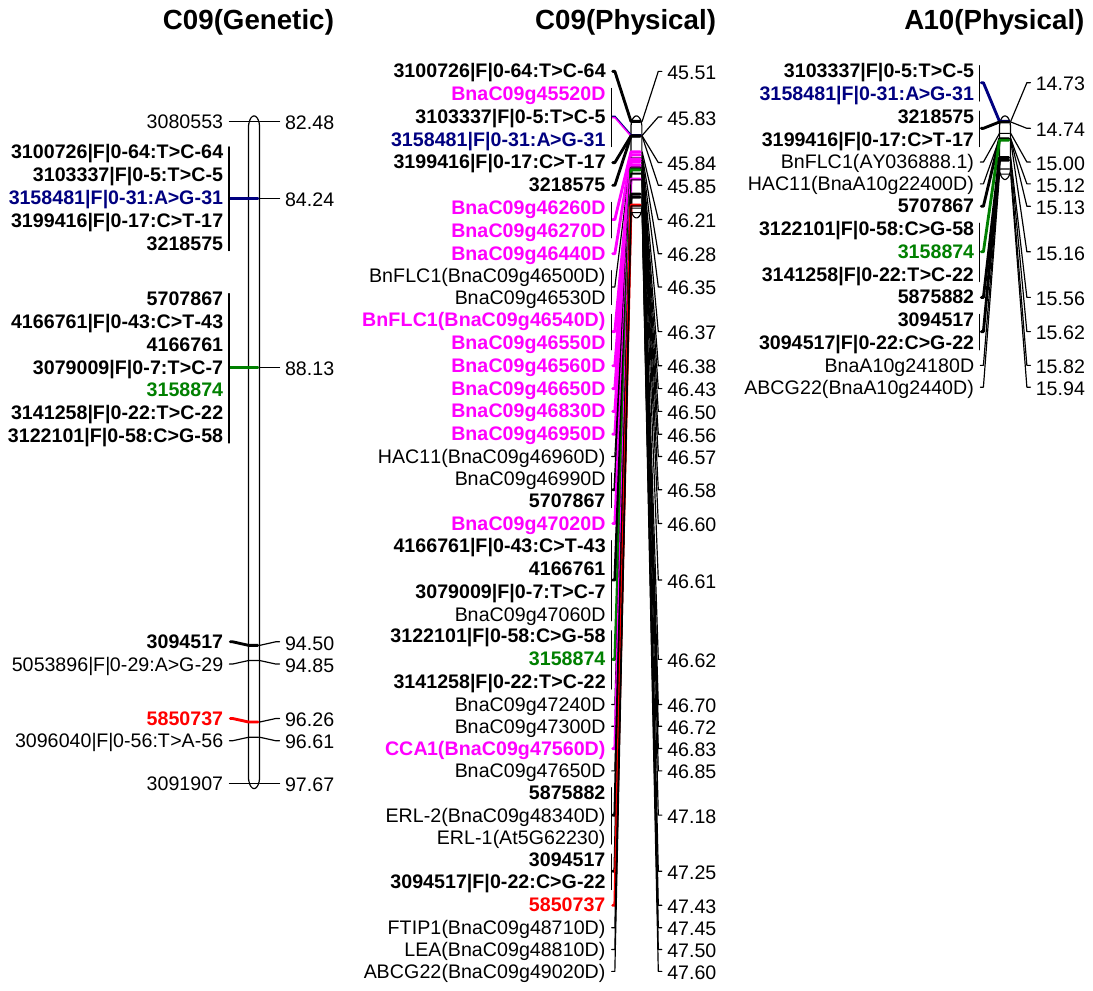
b

Plant height

Δ^13^C

DTF

Δ^13^C

DTF

SY
